# Programmable RNA-guided endonucleases are widespread in eukaryotes and their viruses

**DOI:** 10.1101/2023.06.13.544871

**Authors:** Kaiyi Jiang, Justin Lim, Samantha Sgrizzi, Michael Trinh, Alisan Kayabolen, Natalya Yutin, Eugene V. Koonin, Omar O. Abudayyeh, Jonathan S. Gootenberg

## Abstract

TnpB proteins are RNA-guided nucleases that are broadly associated with IS200/605 family transposons in prokaryotes. TnpB homologs, named Fanzors, have been detected in genomes of some eukaryotes and large viruses, but their activity and functions in eukaryotes remain unknown. We searched genomes of diverse eukaryotes and their viruses for TnpB homologs and identified numerous putative RNA-guided nucleases that are often associated with various transposases, suggesting they are encoded in mobile genetic elements. Reconstruction of the evolution of these nucleases, which we rename Horizontally-transferred Eukaryotic RNA-guided Mobile Element Systems (HERMES), revealed multiple acquisitions of TnpBs by eukaryotes and subsequent diversification. In their adaptation and spread in eukaryotes, HERMES proteins acquired nuclear localization signals, and genes captured introns, indicating extensive, long term adaptation to functioning in eukaryotic cells. Biochemical and cellular evidence show that HERMES employ non-coding RNAs encoded adjacent to the nuclease for RNA-guided cleavage of double-stranded DNA. HERMES nucleases contain a re-arranged catalytic site of the RuvC domain, similar to a distinct subset of TnpBs, and lack collateral cleavage activity. We demonstrate that HERMES can be harnessed for genome editing in human cells, highlighting the potential of these widespread eukaryotic RNA-guided nucleases for biotechnology applications.

## Introduction

Prokaryotic and eukaryotic genomes are replete with diverse transposons, a broad class of mobile genetic elements (MGE). Transposons of the highly abundant IS200/605 family encode a pair of genes: TnpA, which codes for a DDE class transposase responsible for single-strand ‘peel and paste’ transposition, and TnpB, the role of which in the transposon life cycle remain uncertain (*1, 2*). TnpB contains a RuvC-like nuclease domain (RNase H fold) that is related to nuclease domain of the type V CRISPR effector Cas12(*3, 4*), specifically, Cas12 (*5*), suggesting a direct evolutionary path from TnpB to Cas12 (*6*–*8*). This relationship is supported by phylogenetic analysis of the RuvC-like domains, which indicates independent origins of Cas12s of different type V subtypes from distinct groups of TnpBs (*8, 9*). Similarly to the IscB, IsrB, and IshB nucleases, TnpBs are components of obligate mobile element–guided activity (OMEGA) systems, which encode the guide ωRNA adjacent to the nuclease gene, often overlapping the coding region. Biochemical and cellular experiments demonstrated that the ωRNA-TnpB complex is indeed an RNA-guided, programmable DNA endonuclease (*6, 8*).

RuvC domain-containing proteins are not limited to prokaryotes: a set of TnpB homologs, Fanzors, are present in eukaryotes (*7*). Mirroring the diversity of TnpBs in bacteria and archaea, Fanzors have been identified in diverse eukaryotic lineages, including metazoans, fungi, algae, amorphea, and some large double-stranded (ds)DNA viruses. The identified Fanzors fall into two major groups: 1) Fanzor1 proteins are associated with eukaryotic transposons, including Mariners, IS4-like elements, Sola, Helitron, and MuDr, and occur predominantly in diverse eukaryotes; 2) Fanzor2 proteins are found in IS607-like transposons and are present in dsDNA viral genomes. Despite the similarities between TnpB and Fanzors, Fanzors have not been surveyed comprehensively throughout eukaryotic diversity, and have not been characterized experimentally.

Here, we report a comprehensive census of RNA-guided nucleases in eukaryotic and viral genomes, discovering a broad class of nucleases which we named Horizontally-transferred Eukaryotic RNA-guided Mobile Element Systems (HERMES). We examine the diversity of HERMES in eukaryotes, perform a phylogenetic analysis to trace their evolution from prokaryotic ancestors and demonstrate their programmable, RNA-guided endonuclease activity biochemically and in cells.

## Results

### HERMES nucleases are TnpB homologs widespread in eukaryotes and viruses

We identified putative RNA-guided nucleases throughout eukaryotic and viral genomes by comprehensively mining 22,497 eukaryotic and viral assemblies from NCBI GenBank. Our search, seeded with a multiple alignment of RuvC domains from the previously identified Fanzor1 and Fanzor2 proteins(*7*), yielded 3,655 putative nucleases occurring across metazoans, fungi, choanoflagellida, algae, rhodophyta, diverse unicellular eukaryotes, and multiple viral families (Fig. 1A), expanding the known diversity of eukaryotic RuvC homologs, including the Fanzors, about 100-fold (Fig. 1A). Many eukaryotic genomes contain multiple copies of these putative nucleases, suggestive of intragenomic mobility, similar to TnpBs (fig. S1A). Thus, we named these proteins Horizontally-transferred Eukaryotic RNA-guided Mobile Element Systems (HERMES).

**Figure 1:**
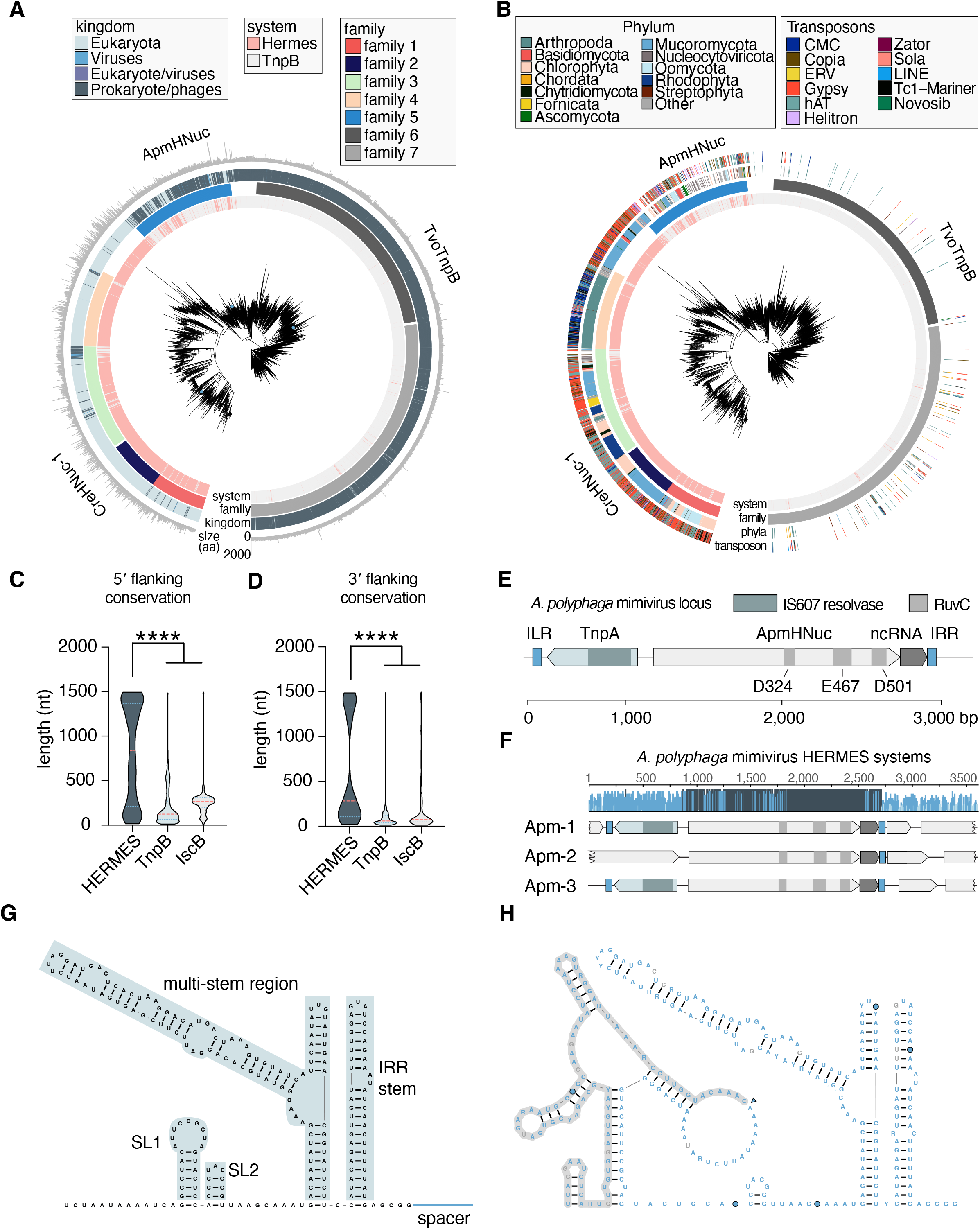
Evolution of HERMES nucleases and their association with non-coding hRNA. A) Phylogenetic tree of representative HERMES and TnpB proteins with the host genome kingdom and HERMES family designation colored. Seven major, strongly supported clades (families 1-7) are color-coded. HERMES and TnpB experimentally studied in this work are labeled. B) Phylogenetic tree of representative HERMES and TnpB proteins with the phyla of their host species and predicted associated transposons marked as rings (all nodes have bootstrap support >0.70). Family and kingdom colors correspond to those in 1A. C) Comparison of predicted ncRNA lengths at the 5′ end of MGE of IscB, TnpB and HERMES systems (****, p<0.0001, one way ANOVA). D) Comparison of predicted ncRNA lengths at the 3′ end of MGE of IscB, TnpB and HERMES systems (****, p<0.0001, one way ANOVA). E) Schematic of the *Acanthamoeba Polyphaga* mimivirus (ApmHNuc HERMES) system, including the HERMES ORF, associated IS607 TnpA, the non-coding RNA region, and the left and right inverted repeat elements (ILR and IRR). F) Conservation of the three HERMES loci in the *Acanthamoeba Polyphaga* mimivirus genome, showing high conservation of the HERMES protein-coding regions and the nearby non-coding RNA. G) Secondary structure of the observed non-coding RNA species from part F, showing significant folding of the non-coding RNA. H) Conserved secondary structure of ApmHNuc HERMES’s non-coding RNA with its most similar HERMES systems.

A phylogenetic tree built from a multiple sequence alignment of HERMES and TnpB revealed 7 major clades (hereafter families 1-7) that were strongly supported by bootstrap analysis, with Fanzor1 represented in families 1-4 and Fanzor2 belonging to family 5 (Fig. 1B). Families 1-4 are each broadly represented in diverse eukaryotes, whereas family 5 is enriched in viruses, including *Phycodnaviridae, Ascoviridae*, and *Mimiviridae* (Fig. 1A-B). In families 1-5, HERMES are interspersed with TnpBs, suggesting multiple captures of TnpB during the evolution of eukaryotes, whereas families 6-7 are predominantly TnpB-containing clades (Fig. 1A-B). In addition, there are a number of unaffiliated HERMES that could not be assigned to any family based on phylogeny.

### HERMES nucleases associate with diverse transposons

Given the previous report on the association of Fanzors with different transposons(*7*), we performed a comprehensive eukaryotic transposon search(*10*) within 10 kb of all HERMES sequences (Fig. 1B). This analysis yielded both previously reported transposon families including Mariner, Helitron, and Sola, and previously undetected ones that include both retrotransposons, such as Gypsy and ERV, and DNA transposons, such as hAT and CMC (fig. S1B). Notably, the two most frequent associations are with the retrotransposon Gypsy and the DNA transposon hAT, suggesting that HERMES might contribute to the retention of these transposons in the respective eukaryotic genomes. Transposon association showed a non-random distribution across the HERMES clades: families 1, 2, 3, and the unaffiliated clade most commonly associated with Gypsy, while family 4 preferentially associated with hAT, CMC, and Tc1-mariner transposons (Fig. 1B). Family 5 associated with a mix of transposons, including Gypsy, Helitron and hAT.

Analysis of the associations of HERMES with surrounding proteins revealed numerous instances of transposase domains, including the serine resolvase found in IS607 elements, further demonstrating the inclusion of HERMES in transposons (fig. S1C). HERMES proteins themselves often contain additional domains beyond the RuvC-like nuclease domain; in particular, family 5 HERMES contain a helix-turn-helix (HTH) and family 7 HERMES contain COG0675 transposase domains, further demonstrating the close similarity between these groups of HERMES and TnpBs (fig. S1D).

### HERMES are associated with conserved, structured non-coding RNAs

Given that TnpB and IscB process either the 3′ end or the 5′ end of the transposon RNA into ωRNA and subsequently form a complex with ωRNA that functions as a RNA-guided dsDNA endonuclease(*6, 8, 11*), we searched all HERMES loci for regions that could encode omega-like RNAs. This search revealed conserved non-coding sequences adjacent to the HERMES coding sequence conservation on both the 5′ and 3′ ends that were considerably longer compared to the respective sequences in TnpB and IscB loci (Fig. 1C-D). This strong conservation of non-coding sequences prompted a detailed search for specific structural hallmarks.

The HERMES proteins in family 5, encoded mostly by large and giant viruses of eukaryotes, are more closely related to TnpB than the HERMES members of families 1-4 (Fig. 1A). Because of this close relationship between TnpBs and HERMES, we initially focused on HERMES of family 5 as a likely source of RNA-guided DNA endonucleases. We selected the HERMES from the *Acanthamoeba polyphaga* mimivirus (ApmHNuc) that is encoded within a IS607 transposon the also encodes a TnpA transposase and contains defined inverted terminal repeats (Fig. 1E).

The *A. polyphaga* mimivirus genome contains three IS607 copies. Alignment of these three loci shows strong sequence conservation with the surrounding HERMES loci to identify conservation throughout the locus (Fig. 1F), we observed high conservation not only within the protein-coding regions but also in the non-coding region at the 3′ ends of the IS607 MGE (Fig. 1E-F), similar to bacterial TnpB. This non-coding sequence conservation extended 200 base pairs (bp) past the end of ApmHNuc ORF, ending upstream of the right inverted repeat (IRR) of the MGE (Fig. 1F). *In silico* RNA secondary structure analysis for the region between the end of the ApmHNuc ORF and the IRR predicted a stable fold (Fig. 1G), suggesting that the transcript of this conserved region could function as a nuclease-associated guide RNA, which we accordingly named HERMES RNA (hRNA). We hypothesized that hRNA forms a complex with ApmHNuc and directs binding and DNA cleavage to a specific sequence in the target. Within the ApmHNuc HERMES cluster, the predicted hRNA structure was highly conserved, and this conservation extended upstream into the coding region of ApmHNuc, indicating possible co-folding with this portion of the coding region and a potential RNA processing site (Fig. 1G). This apparent RNA structure conservation is reminiscent of the OMEGA families, where both the IscB and TnpB families show limited structural variation(*8*), and processing of the upstream region of the co-transcribed mRNA-ωRNA releases functional guide RNAs(*11*).

### ApmHNuc is a hRNA-guided DNA endonuclease

To investigate potential hRNA-ApmHNuc binding, we co-expressed the *A. polyphaga* mimivirus HERMES locus, containing the non-coding RNA region, and *E. coli* codon-optimized ApmHNuc, in *E. coli* (Fig. 2A). Notably, ApmHNuc was unstable when expressed alone and required co-expression with the hRNA for protein stabilization and accumulation (fig. S2), similar to the instability of TnpB in the absence of ωRNA(*6, 8*). We purified the hRNA-ApmHNuc ribonucleoprotein (RNP) and sequenced the RNA component of the complex. Small RNA sequencing revealed enriched coverage between the 3′ ends of the protein ORF and the IRR, in agreement with the evolutionary conservation across the region (Fig. 2B).

**Figure 2:**
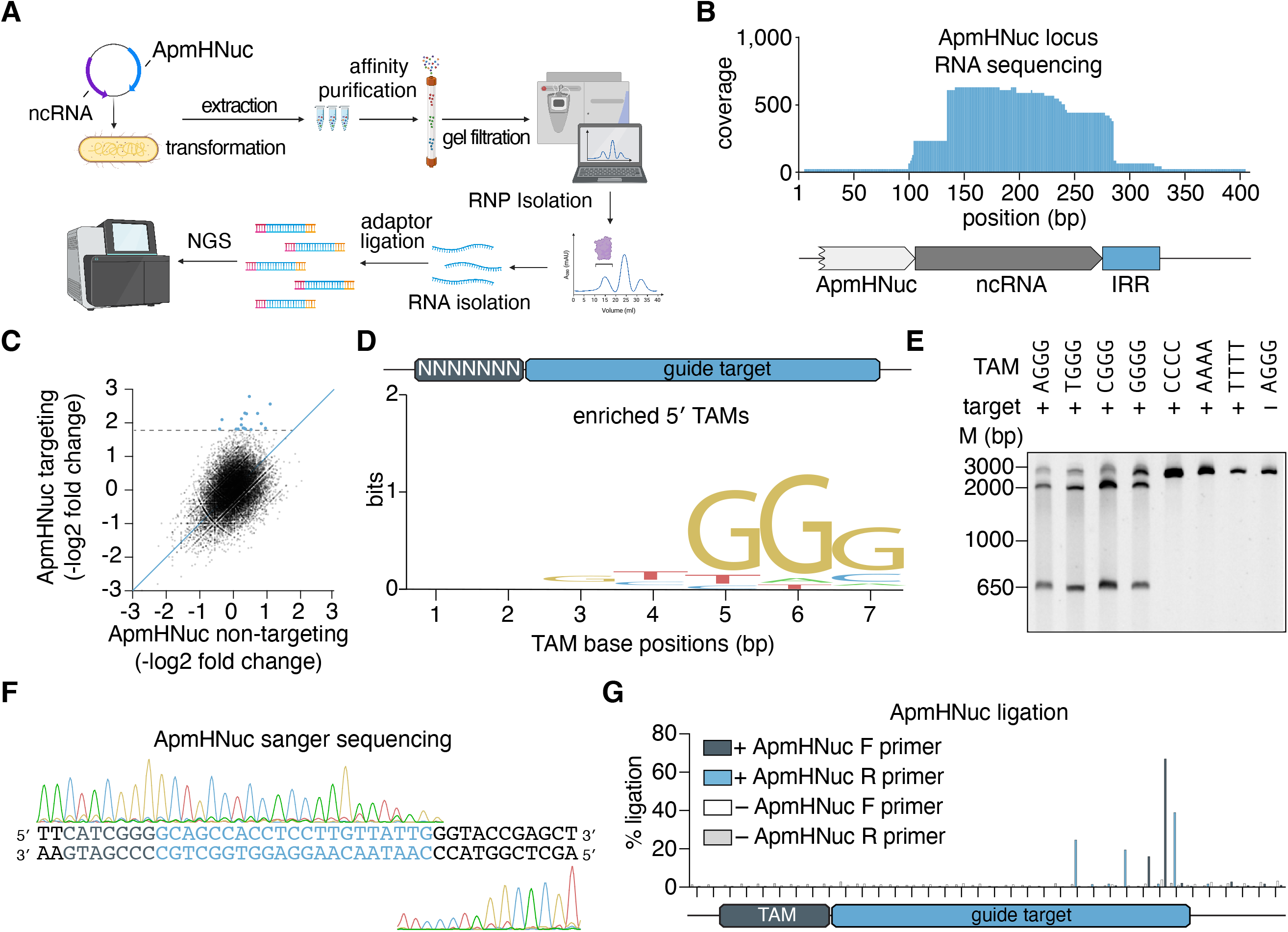
HERMES ribonucleoproteins can be programmed to cleave DNA targets *in vitro*. A) Schematic of the method used for identifying the ApmHNuc associated non-coding RNA. The ApmHNuc protein is co-purified with its non-coding RNA, allowing for the isolation of the non-coding RNA species and identification by small RNA sequencing. B) RNA sequencing coverage of the ApmHNuc-1 non-coding RNA region showing robust expression of the non-coding RNA and its guide sequence extending past the IRR element. C) Scatter plots of the fold change of individual TAM sequences in a 7N library plasmid relative to input plasmid library distribution with either ApmHNuc RNP with a targeting hRNA or a non-targeting hRNA. D) Sequence motif of TAM preference computed from depleted TAMs, showing an NGGG-rich tam preference. E) Biochemical validation of individual ApmHNuc TAM sequences including 4 preferred TAMs (TGGG, AGGG, CGGG, and GGGG) as well as 3 non-TAM sequences and 1 non-targeting sequence. ApmHNuc RNP is incubated with DNA targets containing each of these sequences and cleavage is visualized by gel electrophoresis. F) Sanger sequencing traces of ApmHNuc RNP cleavage on the 5′ CGGG TAM target, showing cleavage downstream of the guide target. G) Next-generation sequencing mapping of the TAM cleavage by ApmHNuc HERMES via NEB adaptor ligation. Cleavage products from *in vitro* cleavage reactions were prepared for sequencing via ligation of sequencing adaptors and PCR prior to next-generation sequencing. Reads were aligned to the TAM target to map cleavage locations. Two separate reactions were ran in parallel with and without addition of ApmHNuc RNP. The cleavage products were amplified in both 5′ and 3′ directions with F denoting 3′ direction and R denoting the 5′ direction.

We hypothesized that ApmHNuc is guided by its associated hRNA to target and cleave DNA sequences. Testing this activity required both the engineering of a reprogrammed hRNA and the determination of sequence preferences, akin to the target adjacent motif (TAM) in the case of TnpB and IscB(*6, 8*). We generated a synthetic hRNA by combining a 3′-terminal 21-nt targeting sequence with the hRNA scaffold determined through RNA profiling. We co-transformed Rosetta cells with plasmids coding for both the synthetic hRNA and ApmHNuc, and isolated the RNP complex from *E. coli*. To determine potential sequence preferences of ApmHNuc, we tested cleavage on a DNA target containing a randomized 7 nucleotide TAM 5′ of a 21 bp target region complementary to the hRNA targeting sequence. We co-incubated this TAM library with purified ApmHNuc RNPs containing either targeting or scrambled synthetic hRNA guide sequences, and profiled the relative depletion of sequences with next-generation sequencing (NGS). The TAM depletion analysis revealed a strong 5′ GGG motif adjacent to the target site (Fig. 2C-D). We validated this TAM on all four possible NGGG sequences, demonstrating robust ApmHNuc cleavage of these sequences, with no detectable cleavage of sequences lacking the TAM (Fig. 2E). In contrast to the G-rich ApmHNuc TAM, TnpB homologs of ApmHNuc universally prefer an A/T rich 5′ TAM (*11*). Interestingly, the GGG motif is present at the start of ApmHNuc MGE sequence and likely contributed to the TAM preference of ApmHNuc.

Cleavage locations of RNA-guided nucleases vary substantially, with cleavage sites located either upstream or downstream of the target sequence. To profile ApmHNuc cleavage patterns, we purified ApmHNuc reaction products and mapped the locations of the cleavage ends using Sanger sequencing. Cleavage occurred in the 3′ regions of the target sequence, with multiple nicks in both the target strand (TS) and the non-target strand (NTS) (Fig. 2F). The cleavage behavior of ApmHNuc at the 3′ end of the target is similar to the cleavage patterns of Cas12 or TnpB nucleases and in general agreement with the properties of programmable RuvC domains(*3, 6, 8*). We sensitively quantified the relative preference for these different nicking sites using an NGS-based assay, finding that during dsDNA cleavage by ApmHNuc the enzyme generated nicks in the NTS at positions 19 and 20, and in the TS at positions 15, 18, and 21 with all cleavage occurring inside the target region, indicating a slightly different cleavage pattern compared to TnpB nucleases (Fig. 2G).

### HERMES nucleases contain a conserved rearranged catalytic site and lack collateral activity

Compared to the majority of the TnpBs, HERMES nucleases contain a substitution in the catalytic RuvC-II motif from a glutamate to a catalytically inert residue (proline or glycine) (Fig. 3A). To determine whether a subset of TnpBs similar to HERMES also contained this substitution, we searched for similarly modified RuvC nuclease domains among the TnpBs. We found a similar apparent inactivation of RuvC-II in TnpBs amongst all seven families (Fig. 3A-B). Given the demonstrated nuclease activity of ApmHNuc, we then searched for conserved acidic residues that could potentially compensate for the RuvC-II-inactivating mutations. Indeed, all HERMES proteins and TnpBs with a loss of the canonical glutamic acid in RuvC-II contained an alternative conserved glutamate approximately 45 residues away (Fig. 3A-B).

**Figure 3:**
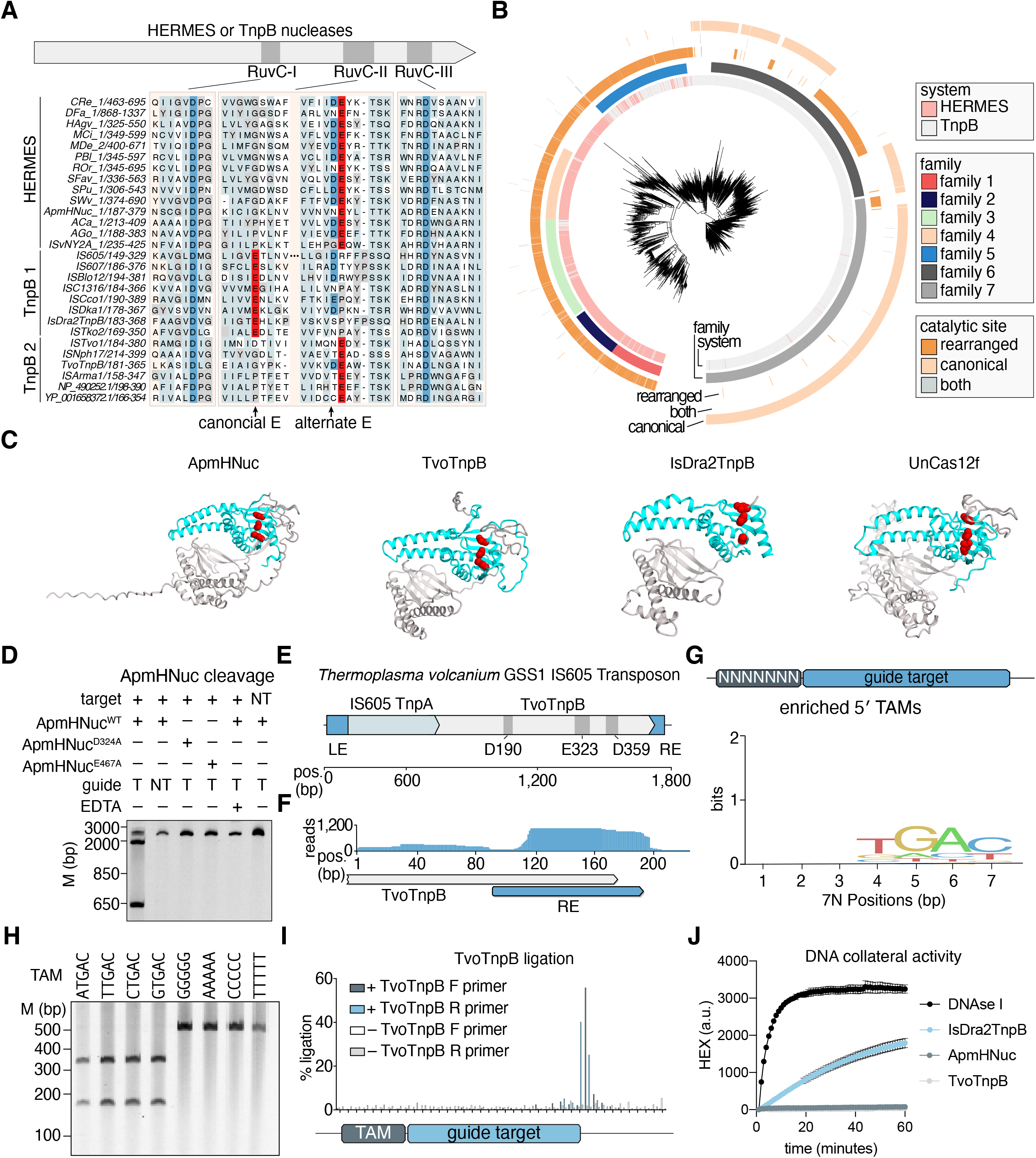
A TnpB with a rearranged catalytic site is also an active nuclease. A) Alignment of the RuvC domains of HERMES and TnpB nucleases showing the alternative glutamate in RuvC-II versus the canonical glutamate. B) Phylogenetic tree of TnpB and HERMES proteins, showing TnpBs and HERMES nucleases with rearranged catalytic sites. C) Predicted AlphaFold-2 structure of ApmHNuc, TvoTnpB, Isdra2TnpB, and Uncas12f, showing that despite having a rearranged glutamate in the RuvC catalytic domain, the catalytic aspartates and glutamates form a putative active catalytic triad (red residues). D) ApmHNuc RNP purified with either targeting (T) or non-targeting (NT) hRNA as well as two catalytic dead ApmHNuc mutants (D324A and E467A) are tested on either a plasmid containing the correct target spacer DNA sequences or a scrambled DNA sequence containing the 5′ TAM TGGG. EDTA is added in lane 5 to quench the cleavage by chelating ions inside the reaction. E) Schematic of the *Thermoplasma volcanium* GSS1TnpB (TvoTnpB) system, including the TnpB with a rearranged catalytic site, associated IS605 TnpA, and the left and right end elements (LE and RE). F) Expression of the non-coding RNA for TvoTnpB, revealing a specific non-coding RNA species that associates with the TvoTnpB protein extending from the ORF to outside the RE element similar to Isdra2TnpB. G) Sequence logo of the TAM for TvoTnpB. H) Biochemical validation of individual TAM preference by TvoTnpB showing that the cleavage by TvoTnpB is TAM (NTGAC) specific. TvoTnpB RNP is incubated with targets containing different 5′ TAMs and cleavage is visualized by gel electrophoresis. I) Next-generation sequencing mapping of the TAM cleavage by TvoTnpB via adaptor ligation. Reads were aligned to the TAM target to map cleavage locations. Two separate reactions were ran in parallel with and without addition of TvoTnpB RNP. The cleavage products were amplified in both 5′ and 3′ directions with F denoting 3′ direction and R denoting the 5′ direction. J) ApmHNuc, TvoTnpB, and Isdra2TnpB DNA collateral cleavage activity are measured using an ssDNA fluorescent reporter, showing a lack of collateral activity for nucleases with the rearranged glutamic acid in RuvC-II. DNase I is used as a positive nuclease control for collateral cleavage activity.

We then compared AlphaFold2-generated structural models of ApmHNuc and a TnpB from *Thermoplasma volcanium* GSS1 (TvoTnpB) that both contain a rearranged catalytic site with the Cryo-EM structures of TnpB from *Deinococcus radiodurans* R1 (Isdra2) and Cas12f from uncultured archaeon (UnCas12f) containing the canonical catalytic site (Fig. 3C) (*12, 13*). This comparison showed that the alternative conserved glutamate of HERMES nucleases and rearranged TnpB (E467 of ApmHNuc and E323 of TvoTnpB) were in close proximity with the catalytic residues in the RuvC-I and RuvC-III motifs, suggesting that these alternative, conserved glutamates compensate for the mutation in RuvC-II (Fig. 3C).

To test the predicted role of the conserved alternative glutamate in HERMES activity, we purified two ApmHNuc RNP mutants at predicted catalytic sites in RuvC-I (D324A) or the alternative glutamate in RuvC-II (E467A) (fig. S3A-C). While the D324A mutant showed no change in the RNP stability during protein purification, we noticed a substantial decrease in the expression of the E467A mutant relative to the wild type protein (fig. S3B). We compared the cleavage efficiencies of these mutants with that of the wild-type ApmHNuc and found, in agreement with the nuclease mechanism, that both RuvC-I and RuvC-II mutants abolished ApmHNuc cleavage activity (Fig. 3D). Thus, the alternative HERMES glutamate is indeed essential for the nuclease activity, which we found was also magnesium dependent (Fig. 3D), similar to other mesophilic RuvC nucleases, and required a temperature range of 30° and 40° C for optimal activity (fig. S3D).

We then evaluated the nuclease activity of TvoTnpB, which also contains the alternative glutamate. We generated TvoTnpB RNPs by co-expressing the TvoTnpB protein with its native locus in *E. coli*, and isolated these RNPs to profile the associated noncoding RNA by NGS. We found significant enrichment of noncoding RNA expression near the right end (RE) element, similar to other TnpB systems (Fig. 3E-F). Applying our TAM assay by coexpressing TvoTnpB with a synthetic ωRNA containing a reprogrammed 21 nt spacer (fig. S4), incubating the RNP with a 7N TAM library plasmid, and sequencing the cleavage products, we found significant enrichment of a TGAC motif near the 5′ target spacer sequence (Fig. 3G). Notably, this TGAC motif is also present at the 5′ end of the left end (LE), marking the beginning of the TvoTnpB-encoding transposon. Because *T. volcanium* is a thermophile, we optimized *in vitro* cleavage efficiency over a range of temperatures and determined the optimal temperature for cleavage of at the TGAC TAM at 60°C (fig. S5A). We validated all four possible NTGAC TAM sequences along with four negative TAM sequences and found TAM-specific cleavage, similar to other HERMES and TnpB nucleases (Fig. 3H). We profiled the ends of the cleavage products with NGS, mapping the cleavage position to position 22 in the non-targeting strand and positions 21 and 22 in the targeting strand (Fig. 3I), with a similar cleavage pattern found by Sanger sequencing (fig. S5B).

We hypothesized that the rearranged RuvC catalytic site of the HERMES might be less solvent exposed, as suggested by the structural analysis (Fig. 3C), reducing acceptance of outside nucleic acids and thus affecting the collateral cleavage activity of the enzyme(*14, 15*). We profiled both ApmHNuc and TvoTnpB for either RNA or DNA collateral cleavage activity by co-incubating the RNP complexes with their cognate targets along with either ssRNA or ssDNA cleavage reporters, single-stranded nucleic acid substrates functionalized with a quencher and fluorophore that become fluorescent upon nucleolytic cleavage. We found that both ApmHNuc and TvoTnpB nucleases indeed lacked detectable collateral DNA and RNA cleavage activity, in contrast to the strong collateral cleavage activity of the canonical TnpB Isdra2TnpB (Fig. 3J and fig. S5C).

### HERMES are widely spread among diverse eukaryotes and are associated with hRNAs

Whereas the Family 5 HERMES that are found primarily in viruses are closely related to TnpBs, most HERMES orthologs, including Fanzor1 nucleases, are more distantly related and have spread through many major branches of eukaryotes, including amoebozoa, fungi, plants, and animals including Chordata and Arthopoda (Fig. 4A). While many HERMES lack introns, as might be expected of TnpB-derived transposons, we also found numerous HERMES with high intron density, up to ∼9.6 introns/kb (Fig. 4B and fig. S6). Intron acquisition supports the notion that HERMES evolved in eukaryotes for extended time.

**Figure 4:**
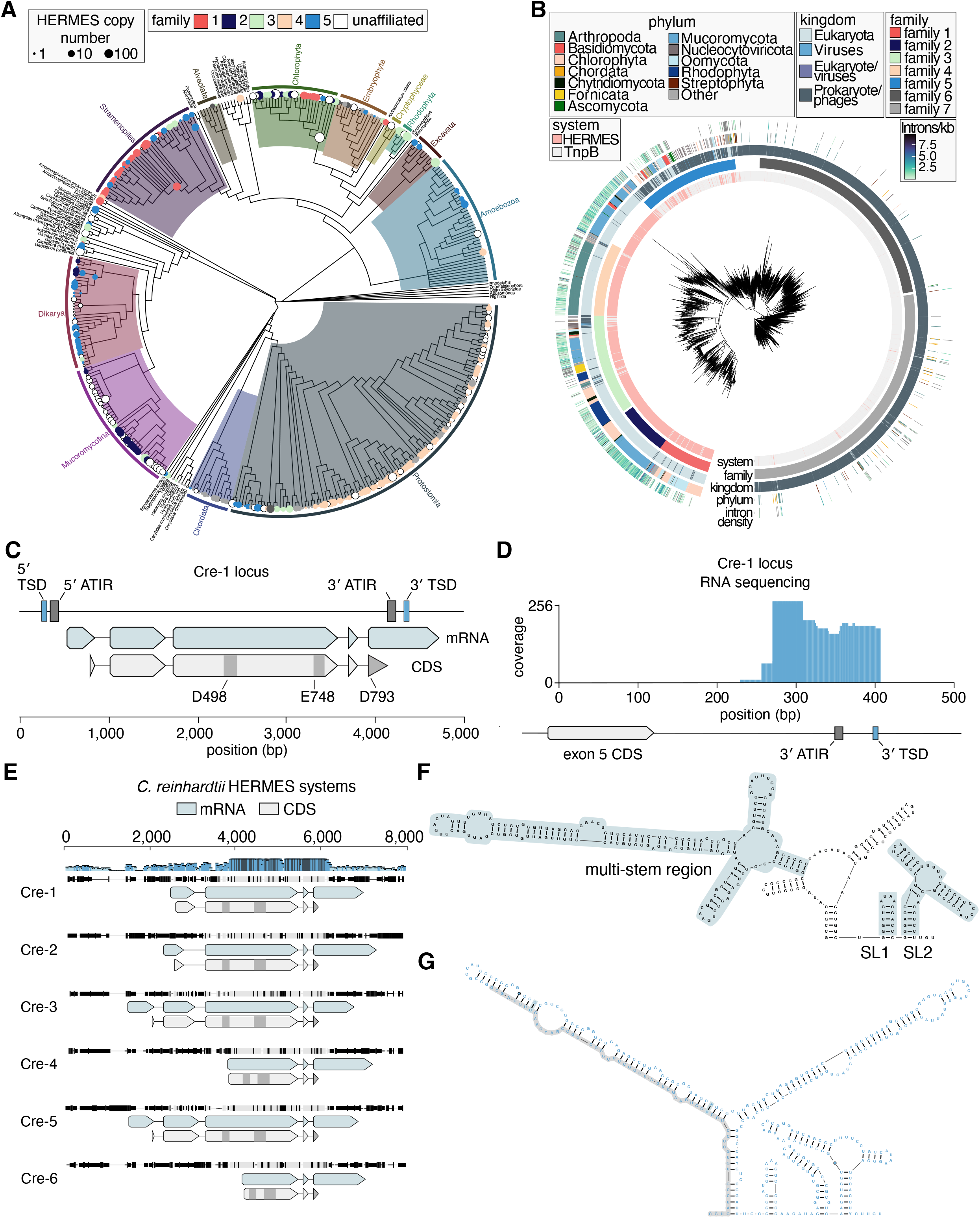
HERMES associated with hRNA are widespread in eukaryotes. A) HERMES systems projected onto the evolutionary tree of eukaryotes (*33*). Nodes and tips of the tree are marked with circles if there are HERMES in the corresponding taxonomic group. Circle sizes are proportional to the HERMES copy number and colored by family. B) Phylogenetic tree of HERMES sequences for which splicing prediction was available. The outer ring shows intron density of the corresponding HERMES genes. C) Schematic of the *Chlamydomonas reinhardtii* HERMES system, including the 5′ asymmetrical terminal inverted repeats (ATIR), 3′ ATIR, 5′ target site duplications (TSD), 3′ TSD, and the mRNA and coding sequences for Cre-1 HERMES. The blue track shows the processed mRNA transcripts relative to the genome and the gray track shows the ORF coding sequences relative to the genome. D) Small RNA sequencing of *Chlamydomonas reinhardtii* showing expression of noncoding RNA at the 3′ end of the CreHNuc that extends beyond the ATIR into the TSD. E) Alignment of all six copies of Cre HERMES inside the annotated part of *Chlamydomonas reinhardtii* genome, showing highly conserved 3′ ends of the Cre HERMES proteins along with its hRNA and variable 5′ end composition of the proteins. The blue track shows the processed mRNA transcripts relative to the genome and the gray track shows the ORF coding sequences relative to the genome. F) Secondary structure of Cre-1 HERMES’ non-coding RNA from 4D-E, showing significant folding of the guide RNA. G) Conserved secondary structure of Cre-1 HERMES’s non-coding RNA and its most similar HERMES systems.

To demonstrate that these diverged HERMES actively process and associate with their cognate hRNAs, we focused on a family 2 HERMES nuclease from the unicellular green alga *Chlamydomonas reinhardtii* (CreHNuc) (Fig. 4C). Using the RuvC profile to search the *C. reinhardtii* genome, we found six RuvC-containing copies of CreHNuc. Notably, these copies contain introns, such that the RuvC domain is encoded in multiple exons. Using RNA sequencing data, we confirmed the presence of four introns within the Cre-1 HERMES pre-mRNA that are spliced out of the mRNA transcript during maturation (Fig. 4C).

The Cre HERMES systems are associated with Helitron 2 transposons, which contain identifiable short target site duplications (TSDs) and asymmetrical terminal inverted repeats (ATIRs). The Cre-1 locus lacks the RepHel domain within the hallmark ATIRs, indicating that it is an non-autonomous Helitron. We hypothesized that either the 3′ TSD or the 3′ ATIR sequence marks the end of the hRNA of Cre-1 and performed small RNA sequencing directly from a *C. reinhardtii* isolate, finding significant enrichment of small non-coding RNAs aligning to the 3′ UTR of the Cre-1 HERMES mRNA (Fig. 4D). The hRNA traces at the Cre-1 locus begin around 100 bp downstream of the end of the last exon and extend across the 3′ ATIR into the TSD (Fig. 4D), suggesting that Cre-1 is involved in the Helitron transposition. We hypothesized that the hRNA of Cre HERMES systems are generally marked by the TSD produced by their native transposon upon insertion. We mapped small RNA-sequencing traces onto all six RuvC-containing copies of CreHNuc and found that all six associated hRNAs lie inside the 3′ UTR of their mRNAs and are strongly conserved between the copies (Fig. 4E and fig. S7A).

Computational secondary structure prediction on the Cre-1 hRNA revealed stable secondary structure, further supporting its potential role in serving as a guide RNA for CreHNuc-1 (Fig. 4F). Generalizing our structural analysis across related CreHNuc genes, we analyzed hRNA conservation within the Cre HERMES clusters. We found that, within the CreHNuc cluster of systems, three representative hRNA structures had high conservation (Fig. 4G), with a conserved upstream region (Fig. 4G, gray region) not present in the RNA-sequencing trace, suggesting possible RNA processing. Moreover, we explored the conservation of this non-coding RNA by searching for similar sequences across the *C. reinhardtii* genome, identifying 20 additional distinct but highly conserved copies of the hRNA (fig. S7B), for a total of 26 copies of Cre HERMES.

We aligned all RuvC-containing Cre HERMES locus from the *C. reinhardtii* genome and observed variable N-terminal compositions (Fig. 4E). Although it remains unclear why the entire coding region of CreHNuc are not conserved similarly to those of ApmHNuc, one possible explanation is that the Helitron transposon undergoes rolling circle replication coupled to transposition that starts at the 3′ end of the transposon, resulting in variable length replicons and truncations.

To evaluate the function of the Cre-1 hRNA, we co-expressed CreHNuc-1 either with its native hRNA on the 3′ end of the MGE or a scramble RNA sequence. We found that CreHNuc is only stable when coexpressed with its hRNA, suggestive of complex formation between the nuclease and hRNA, similar to other HERMES and TnpBs (fig. S7C-D). However, when we co-incubated the RNP with the 7N randomized TAM library plasmids, we did not observe cleavage, suggesting either failure to reconstitute RNP activity in the biochemical context or lack of endonuclease activity in the native CreHNuc.

### HERMES nucleases contain nuclear localization signals and can be adapted for mammalian genome editing

Because eukaryotic RNA-guided endonucleases would need to enter the nucleus to access their genomic targets, we hypothesized that HERMES nucleases might have harbor nuclear localization signals to actively cross the nuclear membrane. In the Alphafold2 predicted structures of ApmHNuc, we identified a disordered region of 64 amino acids at the N-terminus. (Fig. 5A). Computational prediction of the nuclear localization signal (NLS) identified a strong, typical, positive-charged NLS within this N-terminal region of ApmHNuc NLS (fig. S8A).

**Figure 5:**
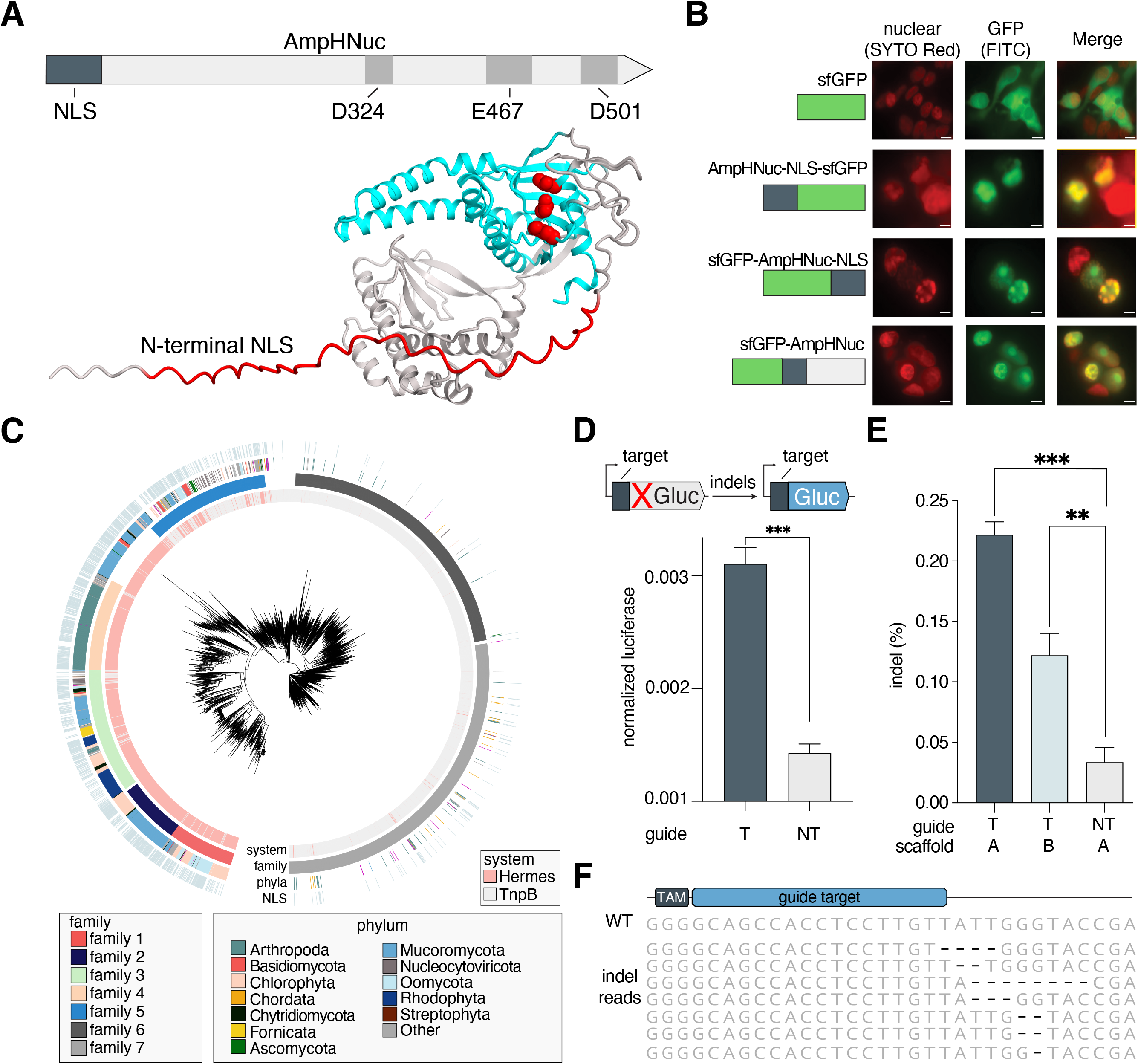
HERMES nucleases contain nuclear localization signals (NLS) and have mammalian genome editing activity. A) Schematic of ApmHNuc HERMES showing the core catalytic triads of split RuvC domain and the predicted N-terminal nuclear localization signal (NLS). The N-terminal NLS like element is colored in red and the catalytic triad is shown as red space filling residues inside the cyan RuvC domain on the AF2 predicted ApmHNuc structure. B) Confocal images of a regular sfGFP, the predicted ApmHNuc NLS fused to sfGFP on either the N-terminal or C-terminal end, and sfGFP fused directly to the N-terminal of ApmHNuc transfected into HEK293FT cells and stained with SYTO Red nuclear stain. Images include the nuclear stain (red), GFP signal (green), and a merged image. Scale bar, 10 μm. C) Phylogenetic tree of HERMES proteins showing which sequences have predicted NLS elements within 15 residues of their N-terminal or C-terminal ends. The phyla and families of the sequences are also marked as rings. D) An ApmHNuc mammalian expression vector and hRNA expression plasmid are co-transfected into HEK293FT cells targeting a luciferase reporter where a Cypridina luciferase (Cluc) is driven by a constitutive promoter and a Gaussia luciferase (Gluc) is placed out of frame from the native start codon. ApmHNuc with a targeting guide against the reporter shows a significantly higher normalized luciferase signal than a non-targeting guide (***, p<0.001, two-sided t-test). E) Indel frequency on the luciferase reporter is measured by next-generation sequencing. The targeting guide with either wild type ApmHNuc hRNA scaffold or T to C mutant scaffold to boost expression is compared against a non-targeting guide. Both scaffolds show a significant increase in indel frequency compared to the non-targeting guide (***, p<0.001, **, p<0.01, one-way ANOVA). F) Representative indel alleles from the targeting guide condition on the luciferase reporter, showing deletions centered around the 3′ end of the guide target.

To evaluate the functionality of the predicted ApmHNuc NLS, we fused the N-terminal portion of ApmHNuc containing the NLS to either the N-terminus or C-terminus of super-folded GFP (sfGFP). We also attached the sfGFP onto the N-terminus of wild-type ApmHNuc and visualized its location via fluorescent microscopy. We found that compared to a wild-type sfGFP, sfGFP with the NLS from ApmHNuc fused to either terminus had strong nuclear localization (Fig. 5B). Fusion of sfGFP with the complete ApmHNuc also caused strong nuclear localization of sfGFP (Fig. 5B). These results suggest that ApmHNuc indeed contains a functional NLS, likely acquired after the capture of the ancestral TnpBs by eukaryotes.

To determine if additional HERMES nucleases might have acquired NLS sequences, we analyzed each HERMES ORF for predicted NLS and found that across all five HERMES families ∼60% of ORFs had readily identifiable NLS, on par with the prediction accuracy of a validated set of NLS-containing proteins (*16*) and substantially greater than NLS predictions for cytosolic human proteins (fig. S8B-C). Thus, it appears likely that a great majority of HERMES nucleases have acquired mechanisms for nuclear import to access the genome and perform their functions.

We tested whether HERMES nucleases could be adopted for mammalian genome editing by codon-optimizing ApmHNuc for mammalian expression and engineering two hRNA guide scaffolds (fig. S9) for expression in mammalian cells. Because the hRNA is longer in length than typical ωRNAs (>350 nt), we co-transfected HEK293T cells with a T7 promoter-driven guide expression plasmid along with human codon-optimized T7 polymerase and wild-type ApmHNuc protein. We designed a reporter plasmid carrying the 21 nt target matching the T7-driven guide in front of a Gaussia luciferase (Gluc) out of frame from the start codon along with a cypridina luciferase (Cluc) driven by a constitutive promoter on the same plasmid to normalize for transfection efficiency. Indel-generating activity would knock the Gluc into frame, allowing for detectable Gluc luciferase activity. Using this reporter system, we found a significant increase in normalized luciferase in the targeting guide condition compared to a non-targeting guide control, suggesting that indels were generated by the ApmHNuc protein (Fig. 5D). We subsequently checked for indels by targeted PCR and NGS and found indel generation in targeting guide conditions for ApmHNuc (Fig. 5E, fig. S9). Finally, we analyzed the indel pattern and found 2-5 bp deletions near the 3′ end of the target site (Fig. 5F), similar to the indel cleavage patterns of other programmable RuvC containing nucleases such as Cas12 or TnpB.

## Discussion

RNA-guided DNA endonucleases are prominent in prokaryotes, with roles in innate immunity by prokaryotic Argonatues(*17*); adaptive immunity by canonical CRISPR systems(*18*–*20*); RNA-guided transposition by CRISPR-associated transposases(*21, 22*), and yet uncharacterized functions in transposon life cycles by OMEGA systems (*6, 8*). In eukaryotes, whereas RNA-guided cleavage of RNA is the cornerstone of the RNA-interference defense machinery and post-transcriptional regulation (*23, 24*), RNA-guided cleavage of genomic DNA has not been demonstrated, to our knowledge. We show here that the previously uncharacterized eukaryotic homologs (*7*) of the OMEGA effector nuclease TnpB are RNA-guided, programmable DNA nucleases. Additionally, we extensively searched diverse genomes of eukaryotes and their viruses to discover thousands of RuvC-containing, which we collectively name HERMES.

Phylogenetic analysis of the HERMES together with TnpBs revealed 7 major families in 4 of which eukaryotic RNA-guided nuclease are mixed with prokaryotic ones, suggesting that TnpB entered the eukaryotic genomes on multiple, independent occasions. Considering the high abundance of TnpBs in bacteria and archaea, and their mobility, along with the exposure of unicellular eukaryotes to bacteria, this apparent history of multiple jumps does not appear surprising. Furthermore, given the wide spread of HERMES in eukaryotes, together with the near ubiquity of TnpBs in bacteria and archaea, it appears likely that TnpBs were originally inherited from both the archaeal and the bacterial partner in the original endosymbiosis that triggered eukaryogenesis (*25*). Subsequent events of TnpB capture by eukaryotes could occur via additional endosymbioses as well as sporadic contacts with bacterial DNA. Notably, however, the high intron density in many HERMES implies their long evolution in many groups of eukaryotes. The history of HERMES family 5, however, is quite distinct. This variety of HERMES that are far more closely similar to TnpB than members of other families likely originated from phagocytosis of TnpB-containing bacteria by amoeba and subsequent spread via amoeba-trophic giant viruses (*26*).

Association of HERMES nucleases with transposases suggests a role for their RNA-guided nuclease activity in transposition similarly to the case of TnpB. The exact nature of that role, however, remains unknown. TnpB has been reported to boost the persistence of the associated transposons in bacterial populations (*27, 28*). TnpB and HERMES potentially could perform different mechanistic roles in transposon maintenance. In particular, these RNA-guided nucleases could target sites from which a transposon was excised, initiating homology directed repair through a transposon-containing locus, restoring the transposon in the original site and thus serving as an alternate mechanism of transposon propagation (*28*). The association of TnpBs and HERMES with diverse types of transposases suggests that the function(s) of the RNA-guided nucleases do not strictly depend on the transposition mechanism.

Our biochemical characterization of the HERMES nucleases revealed both similarities with the homologous TnpB and Cas12 RNA-guided nucleases and several notable distinctions. Similar to TnpB and Cas12, HERMES nucleases generate double-stranded breaks through a single RuvC domain and cleave the target DNA near the 3′ end of the target. However, unlike TnpB and Cas12 enzymes, which have strong collateral activity against free ssDNA, HERMES nucleases and a subset of related TnpBs contain a rearranged catalytic site that is not conducive to collateral activity. In contrast to the T-rich TAMs of TnpB and PAMs of Cas12, the HERMES TAM preference is diverse, with a GC preference observed for Family 5 HERMES (Fanzor2).

Importantly, the TAM preference seems to align with the insertion site sequence, which is compatible with a role of HERMES in transposition. Finally, the hRNA of HERMES overlaps with the transposon IRR and TIR, much like TnpB’s ωRNA, but extends farther downstream of the HERMES ORF, in contrast to the ωRNAs that ends near the 3′ regions of the TnpB ORF. Thus, although the HERMES nucleases originated from TnpB systems, some properties of these eukaryotic RNA-guided nucleases are notably different from those of the prokaryotic ones. Future structural studies will help to elucidate the mechanisms of these biochemical differences.

We also demonstrate that HERMES nucleases can be applied for genome editing with detectable cleavage and indel generation activity in human cells. While the HERMES nucleases are compact (∼500 amino acids), which could facilitate delivery, and their eukaryotic origins might help to reduce the immunogenicity of these nucleases in humans, additional engineering is needed to improve the activity of these systems in human cells, as has been accomplished for other miniature RNA-guided nucleases, such as Cas12f (*29*–*32*). The broad distribution of HERMES nucleases among diverse eukaryotic lineages and associated viruses suggests many more currently unknown RNA-guided systems could exist in eukaryotes, serving as a rich resource for future characterization and development of new biotechnologies.

## Acknowledgments

We would like to thank K.Kato for Alphafold2 structure prediction; D. Weston, and E. Boyden for MiSeq instrumentation support; G. Feng and D. Wang for gel imager support; L. Villiger for guide engineering strategies; K. Chung for FPLC access; Weidong Bao for advice on transposon analysis; and Z. Tang, R. Desimone and J. Crittenden for support and helpful discussions. We thank the members of the Abudayyeh-Gootenberg labs for support and advice.

## Funding

J.S.G. and O.O.A. are supported by NIH grants 1R21-AI149694, R01-EB031957, R01-AG074932, and R56-HG011857; The McGovern Institute Neurotechnology (MINT) program; the K. Lisa Yang and Hock E. Tan Center for Molecular Therapeutics in Neuroscience; G. Harold & Leila Y. Mathers Charitable Foundation; NHGRI Technology Development Coordinating Center Opportunity Fund; MIT John W. Jarve (1978) Seed Fund for Science Innovation; Impetus Grants; Cystic Fibrosis Foundation Pioneer Grant; Google Ventures; FastGrants; Harvey Family Foundation; Winston Fu; and the McGovern Institute. E.V.K is supported through the NIH Intramural Research Program (National Library of Medicine).

## Author contributions

O.O.A., J.S.G, and E.V.K. conceived the study and participated in the design, execution, and analysis of experiments. K.J. designed and performed the experiments in this study, performed computational analyses, and analyzed the data. J.L. developed computational pipelines for HERMES discovery and performed computational analyses. S.S. and M.T. purified proteins and participated in experiments. A.K. assisted with confocal microscopy experiments. N.Y. performed computational analyses related to HERMES discovery, alignment, tree building, and domain discovery. K.J., J.L., E.V.K., O.O.A., and J.S.G wrote the manuscript with help from all authors.

## Competing interests

A patent application has been filed related to this work. J.S.G. and O.O.A. are co-founders of Sherlock Biosciences, Proof Diagnostics, and Tome Biosciences.

## Data and materials availability

Sequencing data will be available at Sequence Read Archive. Expression plasmids are available from Addgene under UBMTA; support information and computational tools are available at https://www.abugootlab.org/.

