## Supplementary Material for "Programmable RNA-guided endonucleases are widespread in eukaryotes and their viruses"

[3]National Center for Biotechnology Information
National Library of Medicine
National Institutes of Health
Bethesda, MD 20894, USA

‡ These authors jointly supervised the work.
*These authors contributed equally.

### Materials and Methods

Computational Discovery of HERMES systems

A profile of the HERMES RuvC domain (HERMES profile) was constructed by aligning the previously discovered Fanzor proteins (seed sequences) with MUSCLE v5 (-align), extracting the RuvC domain, and building a profile HMM with hmmbuild (default options) from the HMMER v3 suite of programs. An initial set of putative HERMES proteins was gathered by searching all annotated proteins and translated ORFs (stop codon to stop codon) longer than 100 residues in NCBI eukaryotic and viral assemblies (one assembly per species) as well as all full length proteins annotated on eukaryotic and viral sequences in GenBank (hmmsearch -E 0.001 -Z 61295632). To predict introns in HERMES ORFs, AUGUSTUS v3.5.0 and Spaln v2.4.13f were applied to the genomic region containing the ORF (10 kb upstream/downstream). AUGUSTUS was used for ab initio gene prediction when there was an available parameter set of the same class as the target species. Tantan was used to soft-mask the genome prior to gene prediction using an "-r" parameter of 0.01 if the genome AT fraction was less than 0.8 and 0.02 otherwise (with the suggested scoring matrix for AT-rich genomes). Spaln was used to splice-align HERMES proteins to the HERMES ORFs (default options). The protein query set for Spaln was generated by searching UniClust90 and GenBank eukaryotic proteins with the HERMES profile. The HERMES profile was iteratively refined by repeatedly searching the initial set of proteins (hmmsearch -E 0.0001 –domE 1000 -Z 69000000), extracting the RuvC domain, clustering with MMseq2 (--min-seq-id 0.5 -c 0.9), aligning the cluster representatives with the profile seed sequences, manually refining the alignment, building a new profile, and using the new profile for the next round. Three rounds of refinement were completed. The refined profile was used for a final round of searches and clusters that would have been included in the profile were kept for the subsequent filtering steps. To reduce the likelihood of including genome assembly contaminants in downstream analysis, all HERMES proteins from NCBI assemblies marked as contig level completeness or those originating from contigs shorter than 50kb (only from assemblies) were discarded. The remaining sequences were clustered using a combination of Diamond v2.1.6 (--evalue 0.0001 --id 70 --query-cover 90 --subject-cover 90 --max-target-seqs 500 --comp-based-stats 3) and MCL (-I 4.0). Each cluster was aligned with MUSCLE and a consensus sequence was computed using a custom python script. The RuvC domains were extracted from each consensus sequence and all aligned with MUSCLE. The alignment was manually inspected and filtered to yield a final set of HERMES sequences.

Computational Discovery of TnpBs

A profile HMM was constructed from a multiple sequence alignment of subsets of HERMES and used to query a custom database of prokaryotic and metagenomic assemblies using HMMER (-E 0.0001 -Z 61295632). Sequences identical to another sequence were discarded and the remaining were clustered with MMseqs2 (--min-seq-id 0.7 -c 0.9 -s 7). The split-RuvC domain was extracted from each cluster representative and further clustered with MMseqs2 (--min-seq-id 0.5 -c 0.9 -s 7) for a two-step clustering process. These split-RuvC domain cluster representatives were aligned with MUSCLE and sequences without alignment to the conserved DED motif were discarded.

Phylogenetic Analysis of HERMES

To make a phylogenetic tree of TnpBs and HERMES sequences, the split-RuvC domain was extracted from every HERMES consensus sequence, clustered with MMseqs2 (--min-seq-id 0.9 -c 0.9) and the cluster representatives were aligned to the split-RuvC domains of the two-step clustered TnpB representatives using MUSCLE (-super5). An approximately-maximum-likelihood phylogenetic tree was constructed with FastTree2 (-lg -gamma) and all branches with a local support value (as computed by FastTree) less than 0.7 were collapsed. The subsequent tree was visualized with R and the ggtree suite of packages.

Prediction of NLS in HERMES

NLStradamus was used with default threshold at 0.6 and model option 2 (four-state bipartite model) to predict NLS domains. For background false positive rate determination, a comprehensive search on Uniprot is performed by looking for homo sapiens cytosolic proteins (with reviewed status) and a total of 1126 proteins are pulled out for analysis. For on target false negative rate determination, the original set of training sequences that include known NLS containing proteins from NLStradamus is used (*16*).

Prediction of HERMES-associated ncRNA

HERMES that were not simply ORF translations were clustered along their entire length at 70% sequence identity and 95% coverage with MMseqs2 (--min-seq-id 0.7 -c 0.95). Each cluster with at least two sequences was subject to ncRNA prediction. For each cluster, the 5' region of the first exon plus 1.5kb upstream bases and 3' region of the last exon plus 1.5kb downstream bases were cut from sequence. The 5' and 3' regions were aligned separately with MAFFT (default options). Each column of the alignment was scored for conservation and the change point in conservation scores was predicted with the R changepoint package to detect a drop in conservation. If the predicted change point was found to be at least 13 bases outside of the exon boundary of every sequence in the alignment, the conserved portion of the exon, plus 11 bases past the change point, were folded with RNAalifold from the ViennaRNA software suite.

HERMES and TnpB Protein Purification

To purify HERMES or TnpB protein, Rosetta2 DE3 pLys cells were transformed with a twin-strep-sumo tag fused to the N-term of a HERMES or TnpB construct along with the predicted hRNA/wRNA driven by a separate vector. Following transformation, single colonies were picked from the agar plate containing antibiotics and picked into a starter culture of 10mL for overnight incubation at 37 degree Celsius. The starter culture was transferred to 2L of TB with the designated antibiotics and grown until the OD reached between 0.6-0.8. The culture was moved to 4C for 30 minutes prior to induction with 0.5mM IPTG induction. The cultures were then grown at 16 degree Celsius overnight and harvested by centrifugation the next day. The pellet is then flash frozen at -80C and subsequently homogenized in lysis buffer (0.02M Tris-HCl pH8.0, 0.5M NaCl, 1mM DTT, and 0.1M cOmplete™, EDTA-free Protease Inhibitor Cocktail (Merck Millipore) with high-pressure sonication for 15 minutes. The homogenized lysates are then centrifuged at 14,000 RPM for 30 minutes at 4C. The clarified supernatant is isolated from the subsequent bacterial pellet and incubated with Strep-Tactin®XT 4Flow® high capacity resin (Cat. No. 2-5030-010) for 1 hour. Following incubation, the crude solution is loaded onto a Glass Econo-Column® Column for gravity flow chromatography and washed three times with the previously described lysis buffer. To elute tagged protein, 10 units of sumo protease is then added onto the column for on-column cleavage overnight at 4C. The next day, the eluent is collected and concentrated through an Amicon® Ultra-15 Centrifugal Filter (Cat. No. UFC9030) before continuing to FPLC. To purify desired protein from added sumo protease, the concentrated eluent is loaded onto a Superdex® 200 Increase 10/300 GL gel filtration column (GE Healthcare). The column was equilibrated with running buffer (10mM HEPES (pH 7.0 at 25C), 1M NaCl, 5mM MgCl2, 2mM DTT). The Peak fractions containing RNP are pulled and analyzed by SDS-PAGE. Correct fractions are concentrated again with amicon filter tubes and subsequently buffer is exchanged into storage buffer (0.02M Tris HCL PH8, 0.25M NaCl, 50% glycerol, 2mM DTT) and stored at -20 for further use. TnpB proteins follow the same purification procedure with the following modifications: T7 express (NEB) pLys strain is used for transformation and subsequent culture.

Small RNA sequencing

Heterologous expression in E. coli: Rosetta2 chemically competent E. coli were transformed with plasmids containing the locus of interest. A single colony was used to seed a 5 mL overnight culture. Following overnight growth, cultures were spun down, resuspended in 750 μL TRI reagent (Zymo) and incubated for 5 min at room temperature. 0.5 mm zirconia/silica beads (BioSpec Products) were added and the culture was vortexed for approximately 1 minute to mechanically lyse cells. 200 μL chloroform (Sigma Aldrich) was then added, culture was inverted gently to mix and incubated at room temperature for 3 min, followed by spinning at 12000xg at 4°C for 15 min. The aqueous phase was used as input for RNA extraction using a Direct-zol RNA miniprep plus kit (Zymo). Extracted RNA was treated with 10 units of DNase I (NEB) for 30 min at 37°C to remove residual DNA and purified again with an RNA Clean & Concentrator-25 kit (Zymo). Ribosomal RNA was removed using a RiboMinus Transcriptome Isolation Kit for bacteria (Thermo Fisher Scientific) as per the manufacturer's protocol using half-volume reactions. The purified sample was then treated with 20 units of T4 polynucleotide kinase (NEB) for 6 h at 37°C and purified again with an RNA Clean & Concentrator-25 (Zymo) kit. The purified RNA was treated with 20 units of 5' RNA phosphatase (Lucigen) for 30 min at 37°C and purified again using an RNA Clean & Concentrator-5 kit (Zymo). Purified RNA was used as input to an NEBNext Small RNA Library Prep for Illumina (NEB) as per the manufacturer's protocol with an extension time of 60 s and 16 cycles in the final PCR. Amplified libraries were gel extracted, quantified by qPCR using a KAPA Library Quantification Kit for Illumina (Roche) on a StepOne Plus machine (Applied Biosystems/Thermo Fisher Scientific) and sequenced on an Illumina NextSeq with Read 1 42 cycles, Read 2 42 cycles and Index 1 6 cycles. Adapters were trimmed using CutAdapt and mapped to loci of interest using BWA-align. Reads were visualized using Genious.

Ribonucleoprotein: RNPs were purified as described. 100 μL concentrated RNP was used as input. The above protocol was followed with the following modifications: 300 μL TRI reagent (Zymo) and 60 μL chloroform (Sigma Aldrich) were used for RNA extraction.

Chlamydomonas reinhardtii was obtained from the University of Minnesota (CRC). The algae was lysed in trizol with glass beads vigorously shaken for 2 hours at room temperature. Then the above protocol was followed with the following modifications: Ribosomal RNA was removed using a plant specific ribominus rRNA depletion kits as per the manufacturer's protocol and the rRNA-depleted sample was purified using Agencourt RNAClean XP beads (Beckman Coulter) prior to T4 PNK treatment. T4 PNK treatment was performed for 1.5 h and purified with an RNA Clean & Concentrator-5 kit (Zymo). Final PCR in the small RNA library prep contained 10 cycles.

<u>Collateral activity testing</u>
DNase alert and RNase alert were purchased from IDT. 1uM of RNP and 10nM of DNA target containing either the target spacer or a scramble spacer are diluted in 1x DNase/RNase alert reaction buffer into 50uL reactions. The solution is mixed well in the reaction test tube and subsequently aliquoted into 384 well plates. The plates are loaded onto applied biosystems qPCR machines and reactions were ran at 37 degree Celcius for ApmHNuc and Isdra2 TnpB, and 60 degree Celcius for TvoTnpB. The SYBR and HEX channel fluorescence intensity is recorded every minute for a duration of 60 minutes. The intensity is normazlied by subtracting the non-target DNA sequence from the target DNA sequence group. A positive control DNAse (2uL) and RNAse (2uL) is ran along with the HEREMES/TnpB group as a positive control to monitor the assay.

<u>Cloning PAM/TAM libraries</u>
Target sequences with 7N degenerate flanking sequences were synthesized by IDT and amplified by PCR with NEBNext High Fidelity 2X Master Mix (NEB). Backbone plasmid was digested with restriction enzymes (pUC19: KPNI and HindIII, Thermo Fisher Scientific) and treated with FastAP alkaline phosphatase (Thermo Fisher Scientific). The amplified library fragment was inserted into the backbone plasmid by Gibson assembly at 50°C for 1 hour using 2X Gibson Assembly Master Mix (NEB) with an 8:1 molar ratio of insert:vector. The Gibson assembly reaction was then isopropanol precipitated by the addition of an equal volume of isopropanol (Sigma Aldrich), the final concentration of 50 mM NaCl, and 1 μL of GlycoBlue nucleic acid co-precipitant (Thermo Fisher Scientific). After a 15 min incubation at room temperature, the solution was spun down at max speed at 4°C for 15 min, then the supernatant was pipetted off and the pelleted DNA has resuspended in 12 μL TE and incubated at 50°C for 10 minutes to dissolve. 2 μL were then transformed by electroporation into Endura Electrocompetent *E. coli* (Lucigen) as per the manufacturer's instructions, recovered by shaking at 37°C for 1 h, then plated across 5 22.7cm x 22.7cm BioAssay plates with the appropriate antibiotic resistance. After 12-16 hours of growth at 37 C, cells were scraped from the plates and midi- or maxi-prepped using a NucleoBond Midi- or Maxi-prep kit (Machery Nagel).

<u>*In vitro* TAM Discovery</u>
1 μM of RNP and 25 ng of TAM library plasmid were incubated at 37 degree for 2 hours in NEB Buffer 3. Reactions were quenched by placing at 4°C or on ice and adding 10 ug RNase A (Qiagen) and 8 units Proteinase K (NEB) each followed by a 5 min incubation at 37°C. DNA was extracted by PCR purification and adaptors were ligated using an NEBNext Ultra II DNA Library Prep Kit for Illumina (NEB) using the NEBNext Adaptor for Illumina (NEB) as per the manufacturer's protocol. Following adaptor ligation, cleaved products were amplified specifically using one primer specific to the TAM library backbone and one primer specific to the NEBNext adaptor with a 12-cycle PCR using NEBNext High Fidelity 2X PCR Master Mix (NEB) with an annealing temperature of 63°C, followed by a second 20-cycle round of PCR to further add the Illumina i5 adaptor. Amplified libraries were gel extracted, quantified by qubit dsDNA kit (Invitrogen) and subject to single-end sequencing on an Illumina MiSeq with Read 1 200 cycles, Index 1 8 cycles and Index 2 8 cycles. TAMs were extracted and visualized by

Weblogo3. Alternatively, a primer set targeting the TAM library plasmid is used to amplify the uncleaved product for 12 cycle and followed by a second 20 cycle rounds of PCR to add the Illumina i5 adaptor. Amplified libraries were gel extracted and subjected to single end sequencing on an Illumina MiSeq with Read 1 200 cycles, Index 1 8 cycles and Index 2 8 cycles. Depletion of TAMs were calculated by comparing to a non-targeting RNP as control and normalized to the original plasmid library distribution.

*In vitro* cleavage assays

Double-stranded DNA (dsDNA) substrates were produced by PCR amplification of pUC19 plasmids containing the target sites and the TAM sequences. All ωRNA and hRNA used in the biochemical assays was *in vitro* transcribed using the HiScribe T7 Quick High Yield RNA Synthesis kit (NEB) from the DNA templates purchased from IDT. Target cleavage assays performed with ApmHNuc contained 10 nM of DNA substrate, 1 μM of protein, and 4 μM of hRNA in a final 1x reaction buffer of NEB Buffer 3. Assays were allowed to proceed at 37°C for 2 hour, then briefly shifted to 50°C for 5 min, and immediately placed on ice to help relax the RNA structure prior to RNA digestion. Reactions were then treated with RNase A (Qiagen), and Proteinase K (NEB), and purified using a PCR cleanup kit (Qiagen). DNA was resolved by gel electrophoresis on Novex 6% TBE polyacrylamide gels (Thermo Fisher Scientific).

Cleavage position mapping by next generation sequencing

1 μM of RNP and 100 ng of the target plasmid were incubated at 37 degree for 3 hours in NEB Buffer 3. Reactions were quenched by placing at 4°C or on ice and adding 10 ug RNase A (Qiagen) and 8 units Proteinase K (NEB) each followed by a 5 min incubation at 37°C. DNA was extracted by PCR purification and adaptors were ligated using an NEBNext Ultra II DNA Library Prep Kit for Illumina (NEB) using the NEBNext Adaptor for Illumina (NEB) as per the manufacturer's protocol. Following adaptor ligation, cleaved products were amplified specifically using one primer specific to the target plasmid (one on 5' sie of the clevage and one on 3' side of the clevage) and one primer specific to the NEBNext adaptor with a 12-cycle PCR using NEBNext High Fidelity 2X PCR Master Mix (NEB) with an annealing temperature of 63°C, followed by a second 20-cycle round of PCR to further add the Illumina i5 adaptor. Amplified libraries were gel extracted, quantified by qubit dsDNA kit (Invitrogen) and subject to single-end sequencing on an Illumina MiSeq with Read 1 100 cycles, Index 1 8 cycles and Index 2 8 cycles.

Mammalian Cell Culture and Transfection

Mammalian cell culture experiments were performed in the HEK293FT line (Thermo Fisher) grown in Dulbecco's Modified Eagle Medium with high glucose, sodium pyruvate, and GlutaMAX (Thermo Fisher), additionally supplemented with 1× penicillin–streptomycin (Thermo Fisher), 10 mM HEPES (Thermo Fisher), and 10% fetal bovine serum (VWR Seradigm). All cells were maintained at confluency below 80%.

All transfections were performed with Lipofectamine 3000 (Thermo Fisher). Cells were plated 16-20 hours prior to transfection to ensure 90% confluency at the time of transfection. For 96-well plates, cells were plated at 20,000 cells/well. For each well on the plate, transfection plasmids were combined with OptiMEM I Reduced Serum Medium (Thermo Fisher) to a total of 10 μL.

Mammalian genome editing

hRNA scaffold backbones were cloned into a pUC19-based human U6 expression backbone and human codon-optimized HERMES proteins were cloned into pCMV-based destination vector by Gibson Assembly. Then 50 ng of protein expression construct, 50 ng of the corresponding guide construct and 20ng of luciferase reporter were transfected in one well of a 96-well plate using lipofectamine 3000 transfection reagent. After 48 hours, reporter DNA was harvested by washing the cells once in 1xDPBS (Sigma Aldrich) and resuspended in 50 µL QuickExtract DNA Extraction Solution (Lucigen) and cycled at 65°C for 15 min, 68°C for 15 min then 95°C for 10 min to lyse cells. 2.5 µL of lysed cells were used as input into each PCR reaction. For library amplification, target reporter regions were amplified with a 12-cycle PCR using NEBNext High Fidelity 2X PCR Master Mix (NEB) with an annealing temperature of 63°C for 15 s, followed by a second 18-cycle round of PCR to add Illumina adapters and barcodes. The libraries were gel extracted and subject to single-end sequencing on an Illumina MiSeq with Read 1 220 cycles, Index 1 8 cycles, Index 2 8 cycles and Read 2 80 cycles. Insertion/deletion (indel) frequency was analyzed using CRISPResso2.

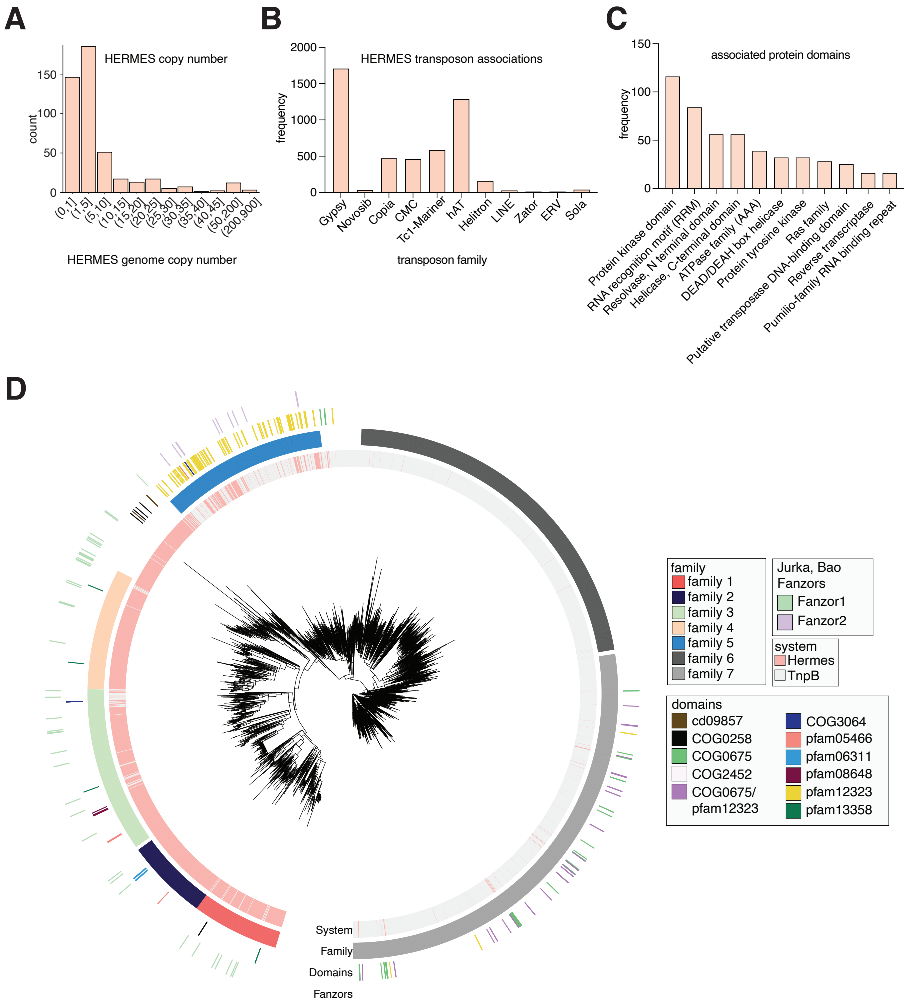


**Fig. S1: Genomic characteristics of HERMES family members**

A) A histogram of the copy number of individual HERMES members inside their respective genomes. B) Frequency of predicted associated transposons nearby HERMES (within +/– 10 kb) per transposon family type. C) Frequency of the top occurring nearby protein domains within 5 genes upstream or downstream of the HERMES MGE. D) Phylogenetic tree of HERMES with the positions of the known Fanzor proteins marked. Phylum and HERMES family information are also marked as rings.

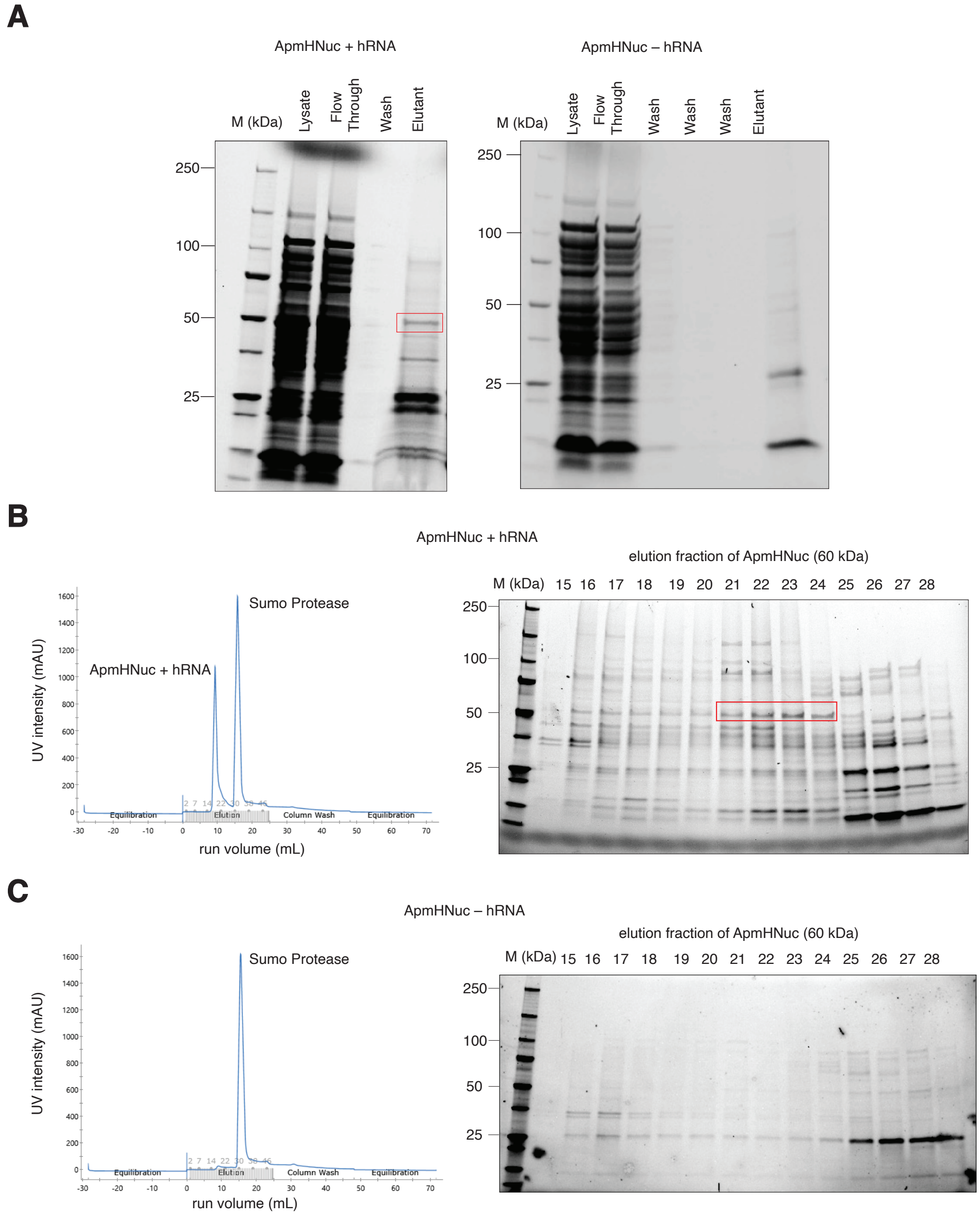


**Fig. S2: Purification of ApmHNuc**

A) Protein gel showing flow through and eluant of AmpHNuc products during gravity flow strep-bead purifications prior to loading of FPLC. Red square denotes the desired protein product. B) FPLC traces of ApmHNuc purified with its hRNA and protein gels showing each fraction's protein products with the desired protein product that was pooled labeled with red squares. C) FPLC traces of AmpHNuc purified without its hRNA and protein gels showing no RNP product in all observed fractions.

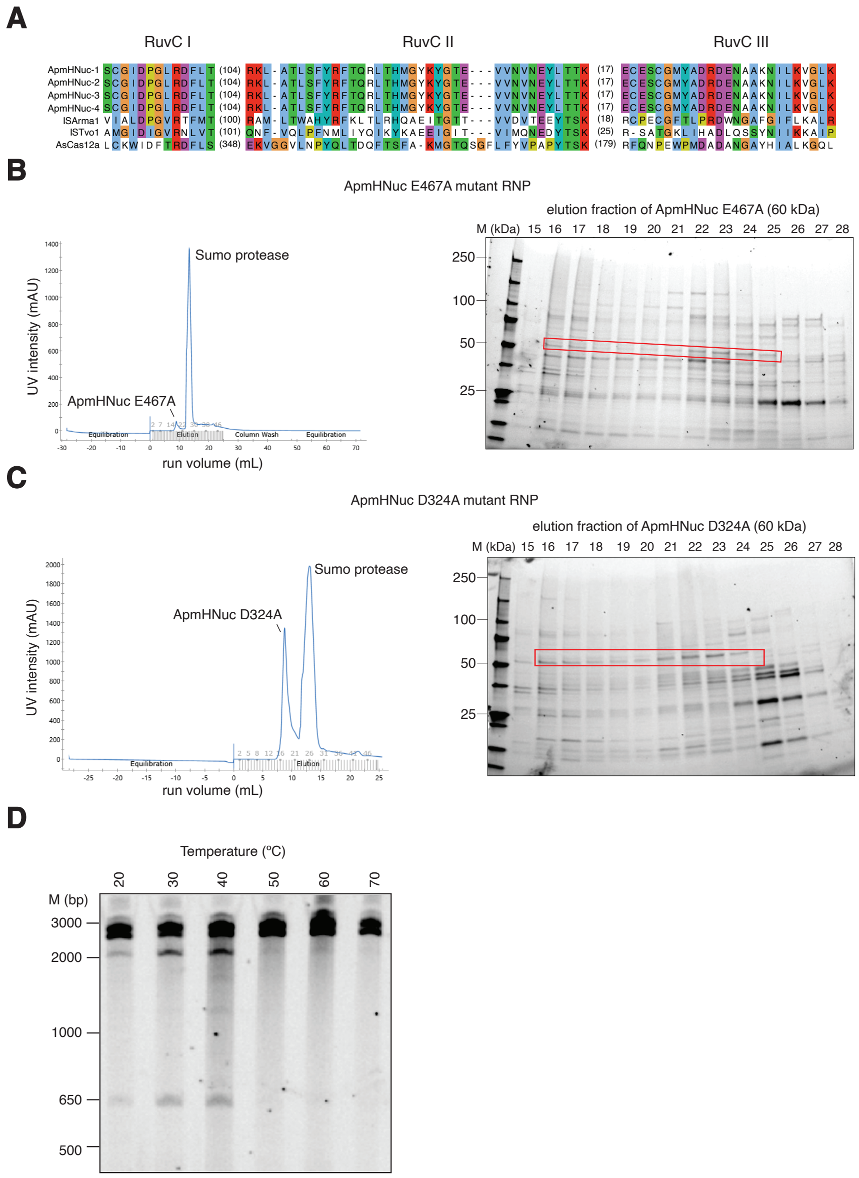


**Fig. S3: Characterization of ApmHNuc nuclease activity**

A) Alignment of ApmHNuc Ruvc domain with Isdra2TnpB RuvC domain to nominate the catalytic RuvC-I aspartic acid (D324) and the RuvC-II glutamic acid (E467A). B) FPLC traces of ApmHNuc E467A mutant purified with its hRNA and protein gels showing each fraction's protein products with the desired protein product that was pooled shown with a red square. C) FPLC traces of ApmHNuc D324A mutant purified with its hRNA and protein gels showing each fraction's protein products with the desired protein product that was pooled shown with a red square. D) Native TBE gel showing nuclease activity of AmpHNuc at temperatures from 10 to 65 degrees Celsius. Reactions were carried out by incubating wild-type ApmHNuc RNP on a plasmid with the TGGG TAM 5′ adjacent to the 21 nt spacer target. Cleavage was visualized by gel electrophoresis.

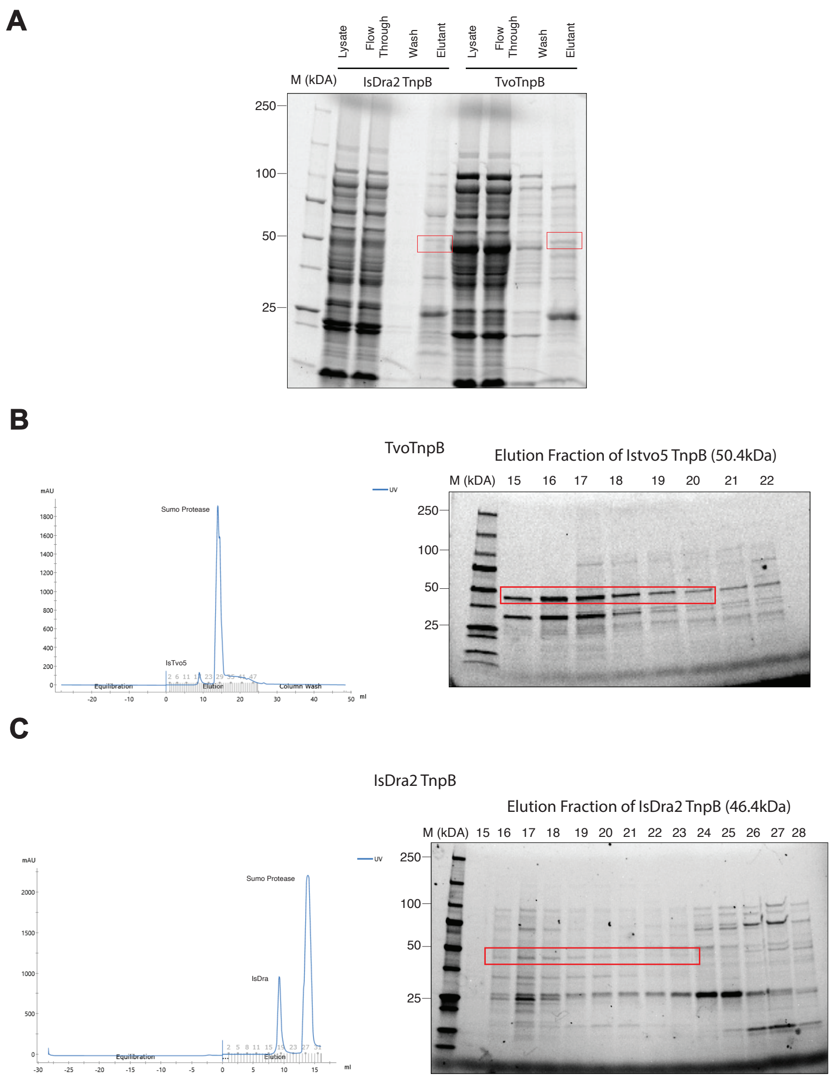


**Fig. S4: Purification of Isdra2TnpB and TvoTnpB**
A) Protein gel showing flow through and eluant fractions of Isdra2TnpB and TvoTnpB products during gravity flow strep-bead purifications. The desired protein product is shown via a red square. B) FPLC traces of TvoTnpB purified with its ωRNA and protein gels showing each fraction's protein products with the desired protein product that was pooled shown with a red square. C) FPLC traces of Isdra2TnpB purified without its ωRNA and protein gels showing each fraction's protein products with the desired protein product that was pooled shown with a red square.

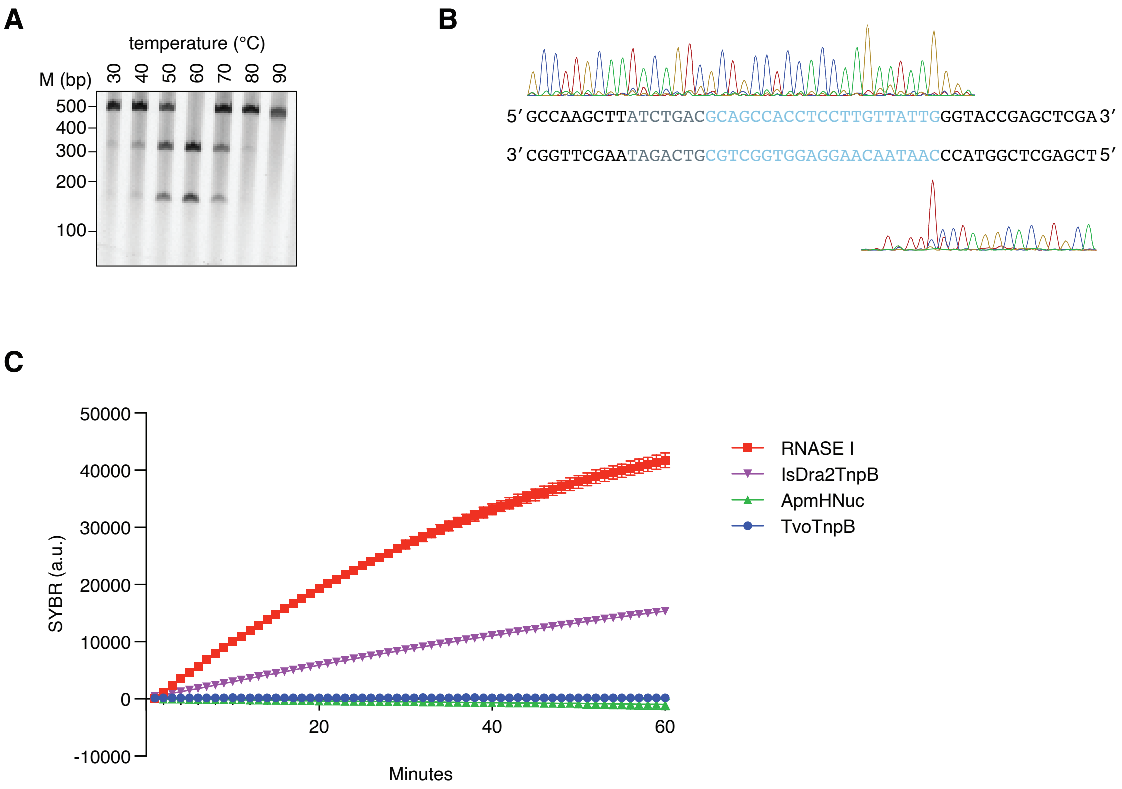


**Fig. S5: Biochemical characterization of TvoTnpB**

A) TvoTnpB DNA cleavage of a 21 nt target containing a 5′ ATGAC TAM at temperatures ranging from 30 degrees Celsius to 90 degrees Celsius, showing optimal cleavage reaction temperature near 60 degrees for TvoTnpB. B) Sanger sequencing traces of TvoTnpB cleavage on a 5′ CTGAC TAM target, showing cleavage at the end of the target. C) Fluorescent signal from RNase alert reporter detection of RNA collateral cleavage activity from RNase A, TvoTnpB, Isdra2TnpB, and ApmHNuc incubated with their target DNA sequences for 1 hour. The signal is normalized to a no DNA target condition.

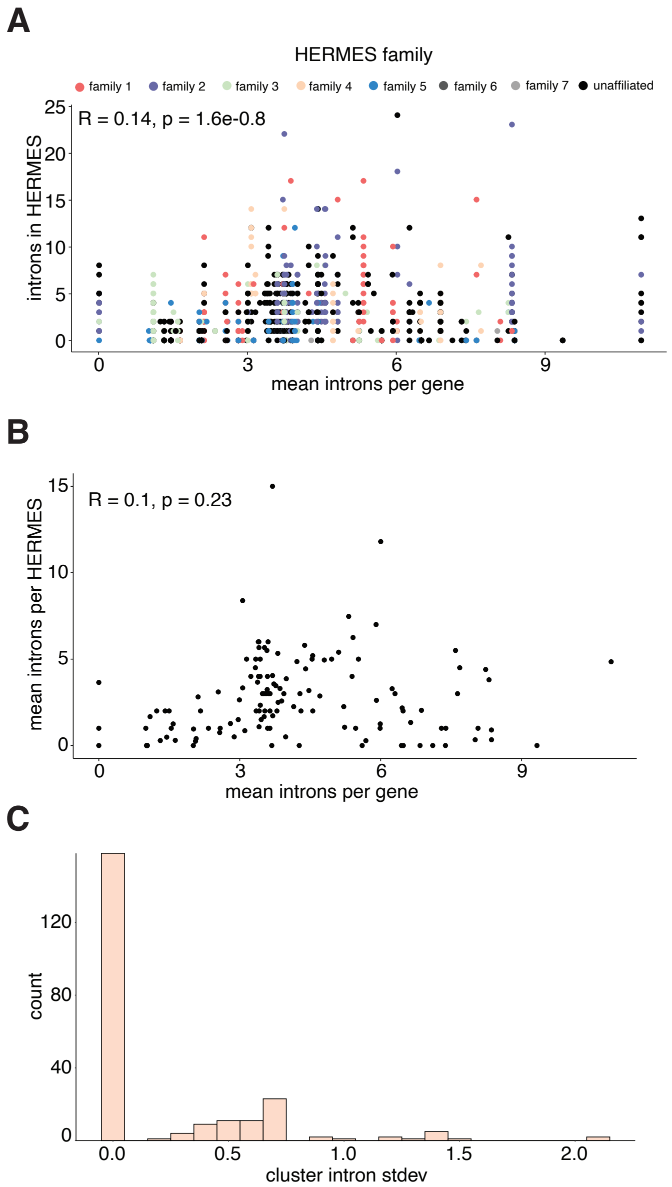


**Fig. S6: Intron characterization of HERMES systems**
A) Comparison of the number of predicted introns in HERMES genes and the mean number of introns per gene in the host genome. Number of introns was defined as the number of exons minus one and calculated from the annotations for the genome provided by GenBank. Correlation and significance values are shown as an inset. B) Comparison of the mean number of introns in HERMES genes in a genome and the mean number of introns per gene in the host genome. Correlation and significance values are shown as an inset. C) Standard deviation of the number of introns per HERMES genes in clusters of 70% sequence identity and 95% alignment coverage. Only sequences with available splicing predictions were clustered and only clusters of two or more sequences are shown.

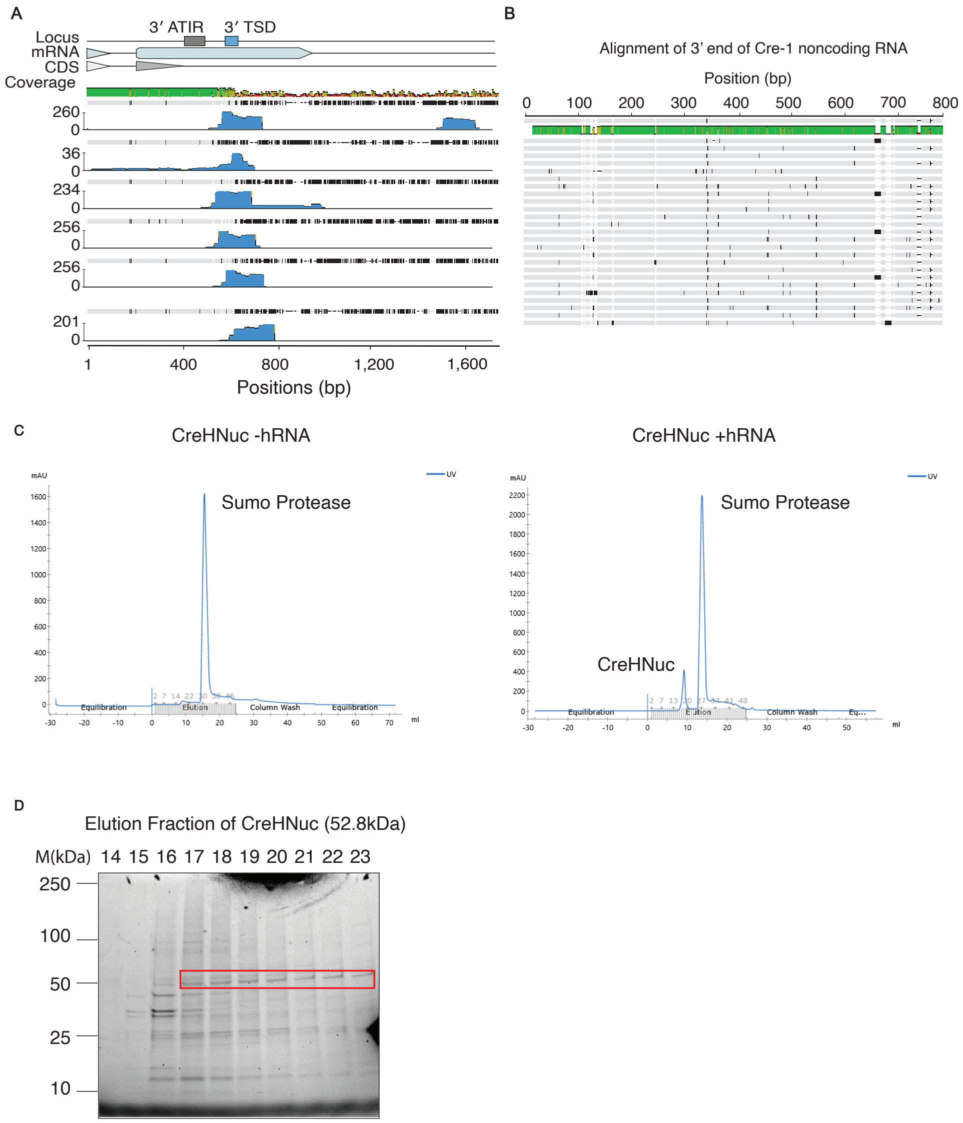


**Fig. S7: Characterization of the CreHNuc hRNAs**
A) Small RNA sequencing traces mapped onto all 6 copies of RuvC-containing HERMES systems in the Cre genome. B) Alignment of the 26 full or partial copies of Cre HERMES MGEs inside the Cre genome at their 3′ end. C) FPLC traces of CreHNuc purified either with or without its hRNA, showing the RNP complex is only stable with the correct hRNA present. The CreHNuc peak in the FPLC trace is labeled. D) Protein gel showing elution fractions of the CreHNuc with the desired protein product that was pooled labeled with a red square.

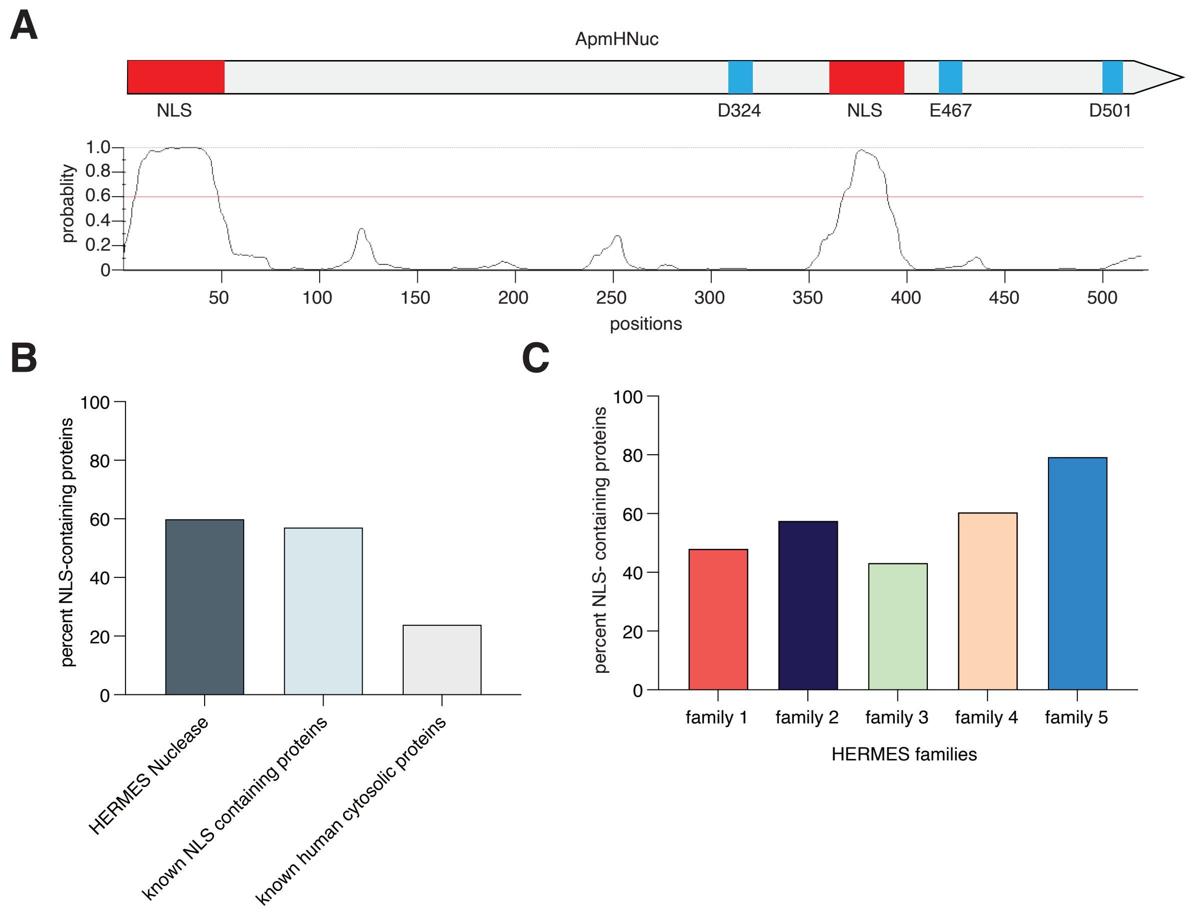


**Fig. S8: ApmHNuc nuclear localization signal characterization**

a) Probability distribution of potential NLS elements across the ApmHNuc protein sequence as predicted by NLStradamus. The default cutoff at 0.6 is used to call significant NLS like elements, revealing one N-terminal NLS and one internal NLS. b) A bar plot of percentage of proteins with predicted NLS on a pool of known human cytosolic proteins (negative control), a set of known NLS containing proteins (positive control) and all HERMES nucleases. c) Per family breakdown of percent NLS continuing HERMES for family 1-5.

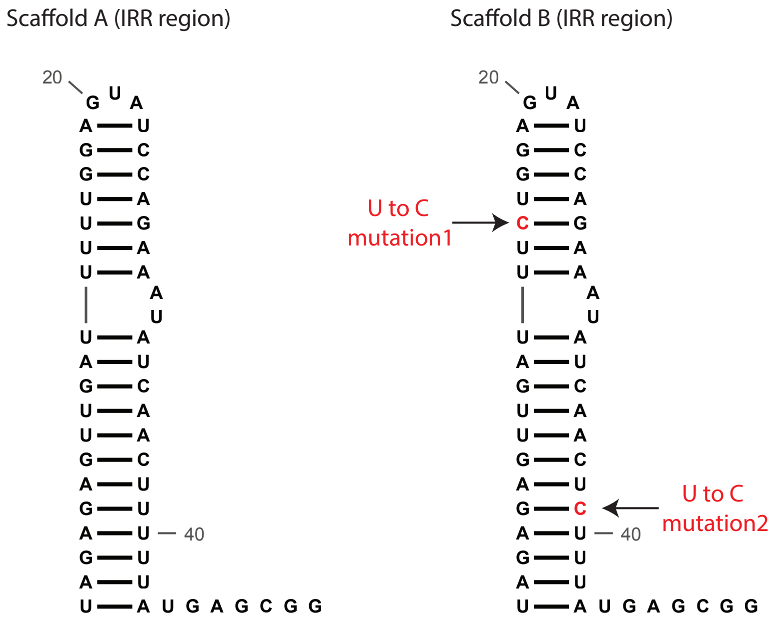


**Fig. S9: Mutating the ApmHNuc hRNA scaffold for small RNA promoter expression**
Mutation of the IRR region in ApmHNuc hRNA which contains two polyU stretches that would terminate small RNA promoter. Two U to C mutations are introduced to delete the polyU stretch in the scaffold B.
